# Effects of Polyphenolic Extracts from Mediterranean Forage Crops on Cholinesterases and Amyloid Aggregation Relevant to Neurodegenerative Diseases

**DOI:** 10.1101/2025.09.28.677303

**Authors:** Antonio D’Errico, Pierluigi Reveglia, Rosarita Nasso, Alessandro Maugeri, Mariorosario Masullo, Rosaria Arcone, Lucia Lecce, Emmanuele De Vendittis, Rosario Rullo

## Abstract

Alzheimer’s and Parkinson’s diseases are the most common neurodegenerative disorders of the central nervous system, characterized by progressive neuronal death and neurological dysfunction. There is currently no treatment that effectively slows the disease progression, and the synthetic drugs proposed to alleviate the symptoms, often cause side effects. Many studies are now focused on neuroprotective properties exhibited by natural agents, such as polyphenols, whose action on different cell signaling pathways is well known. In this article, we analyzed the composition and the properties of polyphenolic extracts from forage plants, such as *Lotus ornitopoidioides*, *Hedysarum coronarium L.*, *Medicago sativa* and *Cichorium intybus*. The LC-MS/MS analysis on the extracts allowed the identification of total 24 phenolic acids and 25 flavonoids. The effects of the extracts on key enzymes of cholinergic neurotransmission, such as acetyl cholinesterase and butyryl cholinesterase, was examined, together with an investigation on the aggregation and disaggregation of amyloid fibrils. The polyphenols acted as inhibitors of the considered enzymes, and interfered with the amyloidogenesis process, with differences depending on the specific extracts. The inhibition constants towards cholinesterases ranged in the 60 – 240 µM interval; the extracts showed different inhibition mechanisms, from competitive to non-competitive. Differences also emerged in the amyloidogenesis process, with IC_50_ values comprised in a large interval. Finally, the extract from *M. sativa* significantly reduced the cell viability of the human neuroblastoma cell line SH-SY5Y. These results suggest that polyphenols extracted from these plants may behave as multitargeting agents against key factors of Alzheimer’s and Parkinson’s diseases. Therefore, they can be considered as promising candidates for the prevention and management of symptoms of these neurodegenerative disorders in combination with pharmacological therapies.

## 1. INTRODUCTION

In elderly people, increased life expectancy is frequently accompanied by the appearance of neurodegenerative disorders of the central nervous system, such as Alzheimer’s disease (AD) and Parkinson’s disease (PD) [1,2]. These pathologies, characterized by a progressive neuronal death, cause severe disabilities, such as memory loss, aphasia, disorientation, depression, tremor, limb rigidity, slowness, walking difficulties, and so on [3]. Furthermore, worsening of these disabilities with age, and the need of long periods for care and therapy, entail high costs even for families of patients affected by these pathologies [4]. AD and PD are the consequence of multifactorial disorders, which alter the neurotransmission metabolism and induce formation of protein aggregates [5,6]. The multifactorial nature of these disorders may explain a missing effective and novel therapeutic treatment, capable to prevent, delay and counteract progression of these diseases [7].

Among enzymes involved in these multifactorial disorders, a central role is played by enzymes involved in the cholinergic neurotransmission, such as acetylcholinesterase (AChE) and butyrylcholinesterase (BuChE) [8,9]. Indeed, these enzymes are primarily responsible for the hydrolysis of acetylcholine or butyrylcholine, respectively, thus allowing return of the activated cholinergic neuron to its resting state [10]. Therefore, AChE and BuChE inhibition by synthetic or natural compounds has relevant implications for the onset of neurodegenerative disorders [7,11,12]. In addition, AChE is also implicated in the aggregation of amyloid-beta (Aβ) peptides leading to the abnormal formation of Aβ plaques around neurons, a hallmark of AD [13]. Therefore, the identification of natural inhibitors of the filamentous aggregation of Aβ peptides would be extremely interesting for a novel therapeutic approach to this neurogenerative disorder [11,14].

Therapies for AD and PD treatment are primarily based on cholinesterase inhibitors, and the most common synthetic drugs, such as donepezil, galantamine and rivastigmine are used against these disorders, although their usage is often accompanied by bothersome side effects, such as loss of appetite, hepatotoxicity and gastrointestinal disorders [15]. To overcome this problem, novel research is focused on the identification of plant-derived natural agents, endowed with neuroprotective properties acting with multi-targeting effects [16–18]. Under these concerns, polyphenols extracted from plants have shown several interesting properties, such as the decrease in the incidence of neurodegenerative diseases, because acting on different cell signaling pathways [19–21]. In particular, flavonoid-rich extracts from Mediterranean plants can modulate AChE and BuChE activity, such as those from Annurca apple flash [18] and lemon peel [22]. In addition, also tannins derived from Mediterranean plants have demonstrated inhibitory activity on AChE and BuChE [23,24]. Flavonoids and tannins, exerting beneficial effects on human health, are present also in plants used as forage crops [25] belonging to the Fabaceae family [26], such as *Lotus ornithopodioides* (known as Southern Bird’s-foot trefoil), *Hedysarum coronarium* (Sulla) [27], *Medicago sativa* (Alfalfa) [28,29], as well as in *Cichorium intybus L.* (Chicory) belonging to the Asteraceae family. All these abundant and edible plants, used in diet and medicine for their strong antioxidant, anti-inflammatory and antimicrobial properties [30,31], represent a suitable source to produce phytochemicals and products for healthcare and treatment of various disorders.

In a recent article, we have described the anti-oxidant properties of polyphenol-or tannin-enriched extracts from leaves of *Lotus ornithopodioides*, *Hedysarum coronarium*, *Medicago sativa*, and *Cichorium intybus* [32]. Now in this study, we focused our attention on the chemical characterization of these extracts and their anticholinesterase activity. To this aim, we determined the effect of these extracts on AchE and BuChE activity, because these enzymes are the major targets for developing new molecules for AD and PD treatment. In addition, the ability of these extracts either to reduce or revert the peptide Aβ_1–40_ self-aggregation was carried out by *in vitro* assays. Finally, in light of potential therapeutic applications of these extracts in prevention, delay and management of AD and PD, we also performed preliminary evaluations of their cytotoxic effect, using *in vitro* cellular models consisting of human neuroblastoma SH-SY5Y cell lines. The results of this investigation show that a natural extract from *M. sativa* may be useful for a possible co-adjutant to a therapeutic approach to AD and PD.

## 2. MATERIALS AND METHODS

### 2.1. Materials and standards

The following reagents MeOH, formic acid, and quinaldic acid were LC-MS grade and purchased from Sigma Aldrich (Darmstadt, Germany). The Phenolic Acids and Alcohols Standard Mixture-V2 along with the Flavonoids Standard Mixture-V2 were obtained from MetaSci library (https://www.metasci.ca/). These substances were used for peak identification, Multiple Reaction Monitoring (MRM) method development, and calibration curves. Acetylthiocholine, butyrylthiocholine, 5’,5’-dithiobis-2-nitrobenzoic (DTNB), thioflavine T, and the enzymes acetylcholinesterase from *Electrophorus electricus* (AChE) and butyrylcholinesterase from equine serum (BuChE), were acquired from Sigma-Aldrich (Milano, Italy). The human β-amyloid peptide (1–40, cat. ab120479) was obtained from Abcam (Cambridge, UK). All other reagents were analytical grade.

### 2.2. Methods

#### 2.2.1. LC-MS/MS analysis of phenolic acids and flavonoids

Plants of Mediterranean forage crops considered in this study were *Lotus ornithopodioides*, *Hedysarum coronarium*, *Medicago sativa*, and *Cichorium intybus*. The origin of these plants and preparation of the relative extracts were extensively reported in our previous work [32]. In particular, extracts from *L. ornithopodioides* and *H. coronarium*, enriched in condensed tannins, were indicated as *Lo*CT and *Hc*CT, respectively, whereas extracts form *M. sativa* and *C. intybus*, enriched in flavonoids, were indicated as *Ms*F and *Ci*F, respectively. The lyophilized material obtained from these extracts was dissolved in dimethylsulfoxide (DMSO) and the different groups of phenolic compounds, such as total phenolics, flavonoids and proanthocyanidins, were determined as previously reported [32]. Total phenolics, measured as gallic acid equivalents, was used for a comparative evaluation of polyphenols among extracts.

The LC-MS/MS system used for the analysis of the extracts included a UHPLC (Nexera Series LC-40, Shimadzu, Kyoto, Japan) coupled to a triple quadrupole/linear ion trap tandem mass spectrometer (QTRAP 4500, AB Sciex, Framingham, MA, USA) that was equipped with a Turbo V ion source. Instrument control, data acquisition, and processing were achieved by the associated Analyst 1.6 and Multiquant 3.0 software. Quantification of phenolic acid and flavonoids was conducted in the same condition as previously reported [33]. The Q1 mass, the Q3 transition, the best parameters and retention times are described in the Supplementary Table S1. Chromatograms for both phenolic acids and flavonoids are reported as Supplementary Figures S1 and S2.

Validation of chromatographic methods was conducted by analyzing calibration curves. Limits of detection (LOD), and quantification (LOQ) are reported in Supplementary Table S2.

#### 2.2.2. Cholinesterase assay and related kinetic parameters

The activity of AChE and BuChE was determined by the Ellman’s method [34], as previously reported [35], using acetylthiocholine or butyrylthiocholine as substrates, respectively. The reaction mixture contained 330 µM DTNB, 500 µM acetylthiocholine or butyrylthiocholine, respectively, and increasing concentrations of the extracts in 500-µL final volume of 0.1 M sodium phosphate buffer, pH 7.4. The reaction started with the addition of 100 mU/mL AChE or BuChE and the absorbance increase at 412 nm was followed kinetically, employing a Cary 100 UV–VIS Spectrophotometer (Agilent, Santa Clara, CA, USA). The steady state enzymatic activity was derived from the initial rate of reaction obtained from the linear part of the kinetics. The inhibitor concentration required to cause 50% reduction of enzymatic activity (IC_50_) was derived from semi-logarithmic plots, in which logarithm of the residual activity ratio was plotted against the extract concentration; the IC_50_ value was extrapolated from the slope of the resulting straight lines [35].

The kinetic parameters *K*_M_ and *V*_max_ of AChE and BuChE activity were determined as previously described [11], by measuring the initial velocity (*v*_i_) of the reaction at different concentrations of the specific thiolated substrate. Values of the inhibition constant (*K*_i_) and the putative mechanism of inhibition were derived comparing the above-mentioned kinetic parameters in the absence or in the presence of fixed concentrations of the extracts, using the following equations for competitive (a), noncompetitive (b) or uncompetitive (c) mechanism:

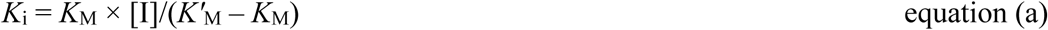

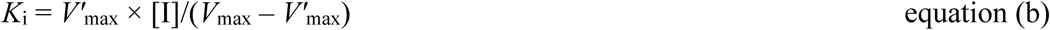

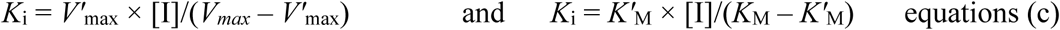

where *K’*_M_ or *V’*_max_ represent the *K*_M_ or *V*_max_ measured in the presence of inhibitor concentration [I].

#### 2.2.3. Assay for self-aggregation or disaggregation of Aβ_1–40_ fibrils

Self-aggregation of Aβ_1–40_ fibrils was achieved as previously reported [11], by incubating for 24 h at 37°C a 12-μL reaction mixture of aggregation buffer, containing 200 mM sodium phosphate buffer, pH 8.0, and 0.5% (v/v) DMSO, in which 96 μM Aβ_1–40_ peptide was dissolved. The reaction ended by adding 0.5 mL of 1.6 μM thioflavine T in 50 mM glycine-NaOH buffer (pH 8.5), and the fluorescence intensity was monitored for 5 min at 25°C, using a Cary Eclipse Spectrofluorimeter (Agilent, Santa Clara, CA, USA). Excitation and emission wavelengths were set at 446 and 490 nm, respectively, and excitation and emission beam slits were both set at 10 nm. The fluorescence value at plateau was averaged over a scan of at least 2 min, and the fluorescence background due to extracts and other components in the reaction mixture was subtracted. To determine the inhibition ratio in the self-aggregation reaction, the decrease of fluorescence signal observed in the presence of various concentrations of extracts was compared to that measured in their absence. The concentration leading to 50% residual Aβ_1–40_ self-aggregation (IC_50_) was derived from a semi logarithmic plot in which the logarithm of the residual activity ratio was plotted against the extract concentration.

The assay for disaggregation of Aβ_1–40_ fibrils entailed a previous fibril aggregation step as reported above. Then, the extent of fibril disaggregation caused by extracts was evaluated by incubating for additional 24 h at 37°C a reaction mixture containing 12 µl of the formed fibril mixture with 2 µl of aggregation buffer, without or with various concentrations of the extracts. The disaggregation ended with the addition of 0.5 mL of the thioflavine T solution and the fluorescence was measured as reported above. The concentration of extract leading to 50% residual Aβ fibrils disaggregation (IC_50_) was derived from a semi logarithmic plot, in which logarithm of the residual activity ratio was plotted against the extract concentration.

### 2.3. Cell cultures and treatments

The human neuroblastoma SH-SY5Y cell line (American Type Culture Collection, Manassas, VA, USA) was maintained in Dulbecco’s modified Eagle medium (DMEM; Microgem Laboratory Research, Milan, Italy), containing 10% heat-inactivated fetal bovine serum (FBS; Microgem Laboratory Research, Milan, Italy), 2 mM L-glutamine, 100 IU/mL penicillin G, and 100 μg/mL streptomycin. Cultures were kept in a humidified incubator at 37 °C with a 5% CO_2_ atmosphere. Cancer cells were subcultured and plated in 75 cm^2^ dishes every two days and were utilized during their exponential phase of growth. Treatments were administered 24 h after plating.

Cell viability was assessed by measuring the mitochondrial metabolic activity, using the 3-(4,5-dimethylthiazol-2-yl)-2,5-biphenyltetrazolium bromide (MTT) assay, as previously reported [18]. Briefly, cells were seeded into 96-well microplates (1 × 10^4^ cells/well). After 24-h incubation, the samples were treated with extracts at varying concentration or with 0.5% DMSO (Sigma-Aldrich, St. Louis, MO, USA) as control vehicle. After 24 h, 10 μL of the MTT solution (5 mg/mL) was added to each well in the dark, and plates were further incubated for 3 h at 37°C under the same culturing conditions. Then, culture medium was removed, and 100 μL of 0.1 N HCl in isopropanol was added to each well to solubilize the formazan crystals. Finally, the absorbance was measured at a wavelength of 570 nm using a BioTek Synergy H1 microplate reader (Agilent, Santa Clara, CA, USA). Cell viability was expressed as a percentage relative to the untreated cells set as 100%.

### 2.4. Statistical analysis

All the assays were performed at least three times and the values obtained were analyzed with the KaleidaGraph program (Synergy, 5.0 version, Adalta, Italy). The kinetic and inhibition parameters were shown as the mean ± standard error. The statistical significance of both nonlinear and linear data fittings was checked using the correlation coefficient R. For the cell viability, statistical significance was determined using ANOVA, followed by Bonferroni’s post hoc test, with significance accepted at p < 0.05.

## 3. RESULTS

### 3.1. Identification of polyphenols extracted from leaves of Mediterranean forage crops

Plants used in this work, namely *Lotus ornithopodioides*, *Hedisarum coronarium*, *Medicago sativa* and *Cichorium intybus L.*, represent a rich source of polyphenols, whose healthy properties as co-adjutants for treatment of neurodegenerative diseases have been described. The relative four extracts, *Lo*CT, *Hc*CT, *Ms*F and *Ci*F, were subjected to LC-MS/MS analysis in order to characterize their constituents, using 81 standard polyphenols, comprising 43 phenolic acids and 38 flavonoids, (Supplementary Table S1). The chromatograms obtained with the four extracts are shown in Fig. 1 and Fig. 2, for phenolic acids and flavonoids, respectively. The results obtained from the MRM analysis are reported in Table 1 and Table 2 for phenolic acids and flavonoids, respectively. Concerning phenolic acids (Table 1), a great variability in composition, concentration, and distribution across the different plants, emerges from the data. Among the 24 identified phenolic acids, several were consistently detected in all extracts, including p-coumaric acid, m-coumaric acid, ferulic acid, 4-hydroxybenzoic acid, phloretic acid, gentisic acid, salicylic acid, and 3,5-dihydroxybenzoic acid, although their concentrations were below the detection limit in some cases. Other phenolic acids, *i.e.* gallic acid, 2,3-dihydroxybenzoic acid, sinapic acid, 2,6-dihydroxybenzoic acid, dihydrocaffeic acid, vanillic acid, and 2,4-dihydroxybenzoic acid were present in *Lo*CT, *Hc*CT and *Ms*F, whereas chlorogenic acid and caffeic acid were present in *Lo*CT, *Hc*CT and *Ci*F. On the other hand, hydroferulic acid and rosmarinic acid were found in *Lo*CT and *Hc*CT, whereas catechol, trans-2-hydroxycinnamic acid, nordihydroguaiaretic acid, and caffeic acid-phenethyl ester were identified only in *Lo*CT. Finally, aspirin was detected and quantified only in *Ms*F. An evaluation of the most abundant phenolic acids quantified in the various extracts was also attempted (Table 1). Indeed, except for phloretic acid, whose concentration was not evaluated, 3,5-dihydroxybenzoic acid, ferulic acid and m-coumaric acid seem to be the most represented compounds in *Lo*CT; gallic acid and vanillic acid in *Hc*CT; salicylic acid, vanillic acid and dihydrocaffeic acid in *Ms*F; chlorogenic acid and caffeic acid in *Ci*F.

**Figure 1.**
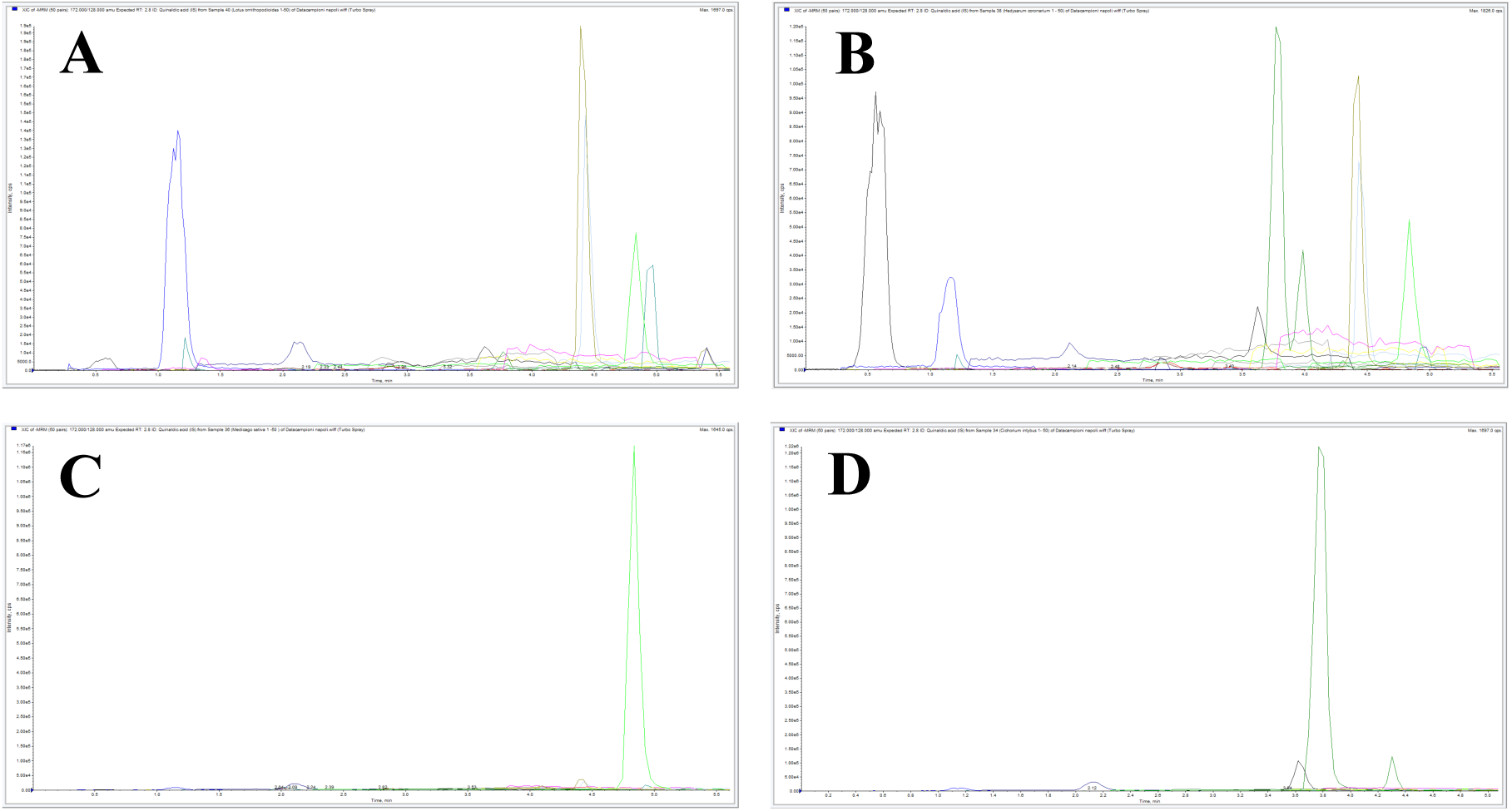
Extract Ion Chromatogram (XIC) of Multi Reaction Monitoring (MRM) of Phenolic Acids extracted from: A) *Lotus ornithopodioides*, B) *Hedysarum coronarium,* C) *Medicago sativa,* D) *Cichorium intybus*.

**Figure 2.**
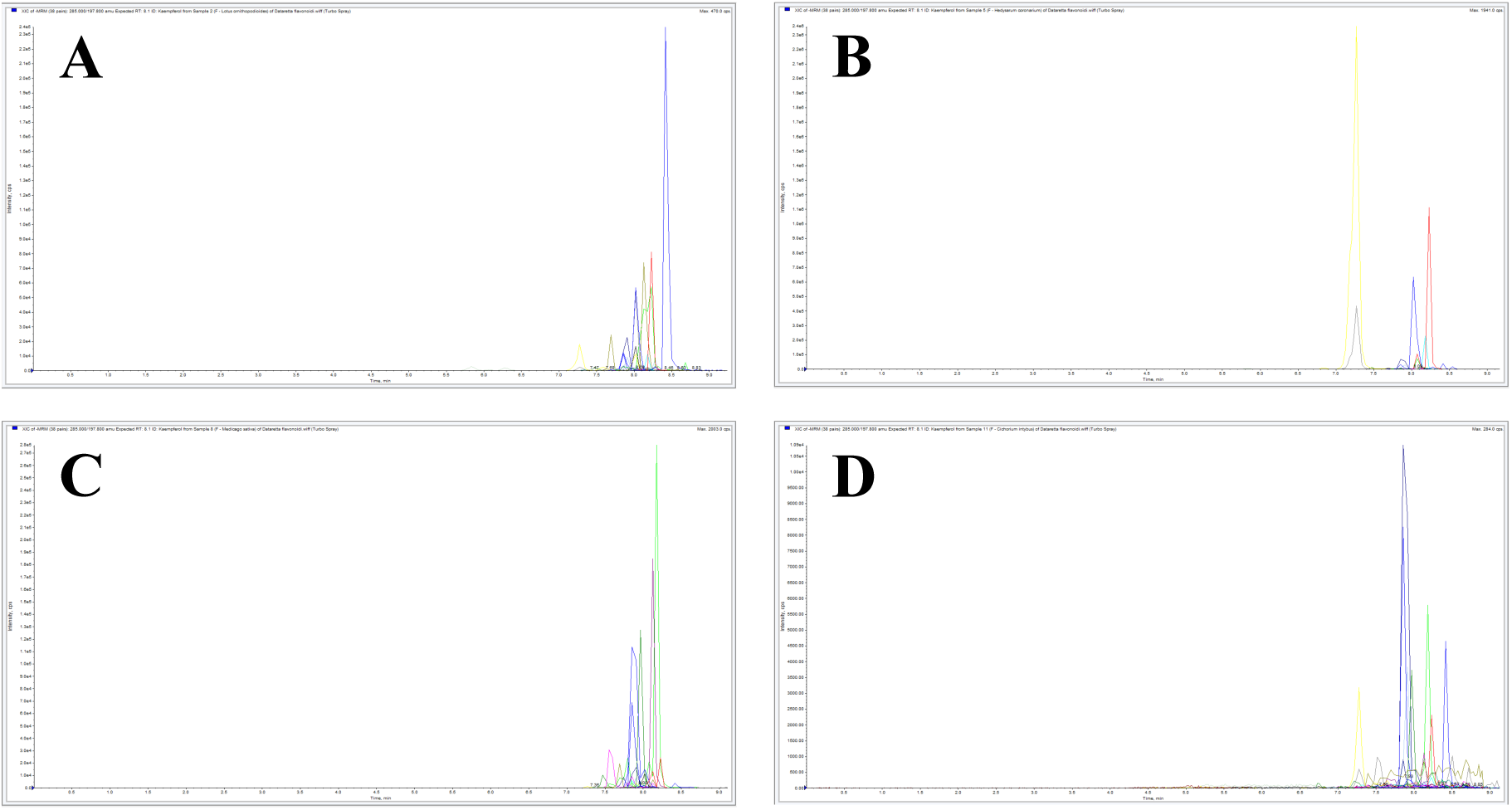
Extract Ion Chromatogram (XIC) of Multi Reaction Monitoring (MRM) of Flavonoids extracted from: A) *Lotus ornithopodioides*, B) *Hedysarum coronarium,* C) *Medicago sativa,* D) *Cichorium intybus*.

**Table 1.** Phenolic acids detected in the extracts of Lotus ornithopodioides, Hedysarum coronarium, Medicago sativa and Cichorium intybus.

| Phenolic acid (µg/mL) | *L. ornithopodioides* | *H. coronarium* | *M. sativa* | *C. intybus* |
| --- | --- | --- | --- | --- |
| Gallic acid | <LOD<sup>a</sup> | 35.10 | <LOD | N/A<sup>b</sup> |
| Chlorogenic acid | <LOD | 9.46 | N/A | 210.18 |
| Caffeic acid | <LOD | <LOD | N/A | 19.65 |
| *p*-Coumaric acid | 1.64 | <LOD | <LOD | <LOD |
| *m*-Coumaric acid | 22.91 | 11.07 | 2.45 | <LOD |
| 2,3-Dihydroxybenzoic acid | <LOD | <LOD | <LOD | N/A |
| Ferulic acid | 24.24 | <LOD | 2.28 | <LOD |
| Sinapic acid | 3.72 | 2.73 | 4.89 | N/A |
| Aspirin | N/A | N/A | 4.15 | N/A |
| 4-Hydroxybenzoic acid | <LOD | <LOD | <LOD | <LOD |
| 2,6-Dihydroxybenzoic acid | <LOD | <LOD | <LOD | N/A |
| Dihydrocaffeic acid | <LOD | <LOD | 20.33 | N/A |
| Phloretic acid | +<sup>c</sup> | + | + | + |
| Hydroferulic acid | <LOD | <LOD | N/A | N/A |
| Catechol | <LOD | N/A | N/A | N/A |
| Gentisic acid | <LOD | <LOD | <LOD | <LOD |
| Salicylic acid | <LOD | <LOD | 108.53 | <LOD |
| trans-2-Hydroxycinnamic acid | <LOD | N/A | N/A | N/A |
| 3,5-Dihydroxybenzoic acid | 63.53 | 5.58 | <LOD | <LOD |
| Vanillic acid | 1.06 | 31.86 | 31.36 | N/A |
| Nordihydroguaiaretic Acid | <LOD | N/A | N/A | N/A |
| Rosmarinic acid | <LOD | <LOD | N/A | N/A |
| Caffeic acid phenethyl ester | <LOD | N/A | N/A | N/A |
| 2,4-Dihydroxybenzoic acid | <LOD | <LOD | <LOD | N/A |

<sup>a</sup> <LOD, below the Limit of Detection.
<sup>b</sup> N/A, not detected.
<sup>c</sup> +, detected in the extract, but non quantifiable in the selected linearity range.

**Table 2.** Flavonoids detected in the extracts of Lotus ornithopodioides, Hedysarum coronarium, Medicago sativa and *Cichorium intybus*.

| Flavonoid (µg/ml) | *L. ornithopodioides* | *H. coronarium* | *M. sativa* | *C. intybus* |
| --- | --- | --- | --- | --- |
| Quercetin | 3.18 ± 0.2 | 28.91 ± 0.5 | 6.7 ± 0.24 | 1.81 ± 0.53 |
| (-)-Epicatechin | N/A<sup>a</sup> | 1.48 ± 0.12 | 13.16 ± 0.7 | N/A |
| (+)-Catechin | 11.17 ± 0.82 | 13.15 ± 1.37 | 24.74 ± 0.93 | 10.39 ± 0.28 |
| Myricetin | <LOQ<sup>b</sup> | <LOQ | <LOQ | N/A |
| Polydatin | N/A | +<sup>c</sup> | N/A | + |
| Kaempferol | 7.74 ± 0.09 | 20.92 ± 3.47 | 26.17 ± 2.43 | 3.99 ± 0.97 |
| Acacetin | N/A | 4.83 ± 0.42 | 4.57 ± 0.52 | N/A |
| Baicalein | 2.04 ± 0.026 | 2.72 ± 0.04 | 2.02 ± 0.03 | 2.12 ± 0.04 |
| (+)-Taxifolin | <LOQ | <LOQ | N/A | N/A |
| Diosmetin | 3.54 ± 0.09 | 2.14 ± 0.4 | 9.54 ± 1 | <LOQ |
| Morin | 6.36 ± 1.33 | 28.42 ± 1.98 | 2.19 ± 0.09 | <LOQ |
| (-)-Epigallocatechin gallate | <LOQ | N/A | N/A | N/A |
| (+/-)-Naringenin | 1.38 ± 0.08 | 0.37 ± 0.03 | 0.83 ± 0.02 | N/A |
| Baicalin | 1.86 ± 0.11 | N/A | 16.01 ± 0.27 | 1.11 ± 0.02 |
| Hesperidin | 23.73 ± 0.08 | N/A | N/A | <LOQ |
| Fisetin | N/A | 2.56 ± 0.03 | N/A | N/A |
| Apigenin | 2.31 ± 0.03 | 1.82 ± 0.04 | 45.42 ± 4.36 | 1.93 ± 0.03 |
| trans-Pterostilbene | 36.33 ± 2.08 | 775.47 ± 70.45 | 2.74 ± 0.05 | N/A |
| Rutin | 4.6 ± 0.79 | 771.23 ± 72.12 | <LOQ | 0.03 ± 0.02 |
| Daidzein | <LOQ | <LOQ | N/A | N/A |
| Hesperetin | 2.4 ± 0.06 | 3.36 ± 0.39 | 1.66 ± 0.02 | 1.47 ± 0.02 |
| Isoliquiritigenin | + | + | + | + |
| Biochanin A | 4.11 ± 0.62 | 13.67 ± 0.17 | 2.64 ± 0.18 | <LOQ |
| Formononetin | 0.69 ± 0.13 | 10.02 ± 0.24 | 0.08 ± 0.03 | <LOQ |
| Genistein | 5.1 ± 0.07 | 7.03 ± 0.17 | 23.48 ± 0.03 | N/A |

<sup>a</sup> N/A, not detected; <sup>b</sup> <LOQ, below the Limit of Quantification; <sup>c</sup> +, detected in the extract, but non quantifiable in the selected linearity range.

Concerning the 25 flavonoids identified in the four extracts (Table 2), also in this case a great variability in their concentration, composition and distribution emerges from the data. In particular, quercetin, (+)-catechin, polydatin, kaempferol, baicalein, diosmetin, morin, apigenin, rutin, hesperetin, isoliquiritigenin, biochanin A, and formononetin were identified in all extracts, although their concentration was below the quantification limit in some extracts. Myricetin, (+/-)-naringenin, trans-pterostilbene, and genistein were present in *Lo*CT, *Hc*CT and *Ms*F, whereas baicalin was present in *Lo*CT, *Ms*F, and *Ci*F. On the other hand, (+)-taxifolin and daidzein were found in extracts from *Lo*CT and *Hc*CT, whereas (–)-epicatechin and acacetin in extracts from *Hc*CT and *Ms*F, and hesperidin in *Lo*CT and *Ci*F. Lastly, fisetin was exclusively detected and quantified in *Hc*CT, and (–)-epigallocatechin gallate was detected, but not quantified in *Lo*CT. An evaluation of the most abundant flavonoids quantified in each extract was also attempted (Table 2). Indeed, with the exception of isoliquiritigenin and polydatin, whose concentrations were not evaluated, trans-pterostilbene and hesperidin seem to be the most represented compounds in *Lo*CT; trans-pterostilbene and rutin in *Hc*CT; apigenin, kaempferol, (+)-catechin and genistein in *Ms*F; (+)-catechin, and kaempferol in *Ci*F.

### 3.2. Effect of the plant extracts on cholinesterase activity

The effects of *Lo*CT, *Hc*CT, *Ms*F and *Ci*F on the steady state activity of cholinesterases was investigated. The dose-dependent inhibition profile exerted by the four extracts on AChE and BuChE activity is shown in Fig. 3. In the AChE assay (Fig. 3A), a significant and progressive reduction of the steady state activity was observed with *Ms*F, whereas the other extracts were much less effective; indeed, when the extracts were added at 100 µM concentration, the residual AChE activity dropped to nearly 20% with *Ms*F, whereas remained above 50% with *Lo*CT, *Hc*CT or *Ci*F. In the BuChE assay (Fig. 3C), the best inhibition profile was observed with *Hc*CT, followed at a distance by *Ms*F, whereas *Lo*CT and *Ci*F were much less effective, even when added at 100 µM concentration. These data were also analysed through a logarithmic transformation of the residual activity ratio. The semi-logarithmic plots drawn for AChE (Fig. 3B) and BuChE (Fig. 3D) allowed the extrapolation of the inhibitor concentration that caused 50% reduction of activity (IC_50_) through the slope of the resulting linearized inhibition profile. The IC_50_ values reported in Table 3 confirm that, among the four extracts, *Ms*F (52 ± 7 µM) and *Hc*CT (40 ± 6 µM) were endowed with the best inhibition power towards AChE and BuChE, respectively.

**Figure 3.**
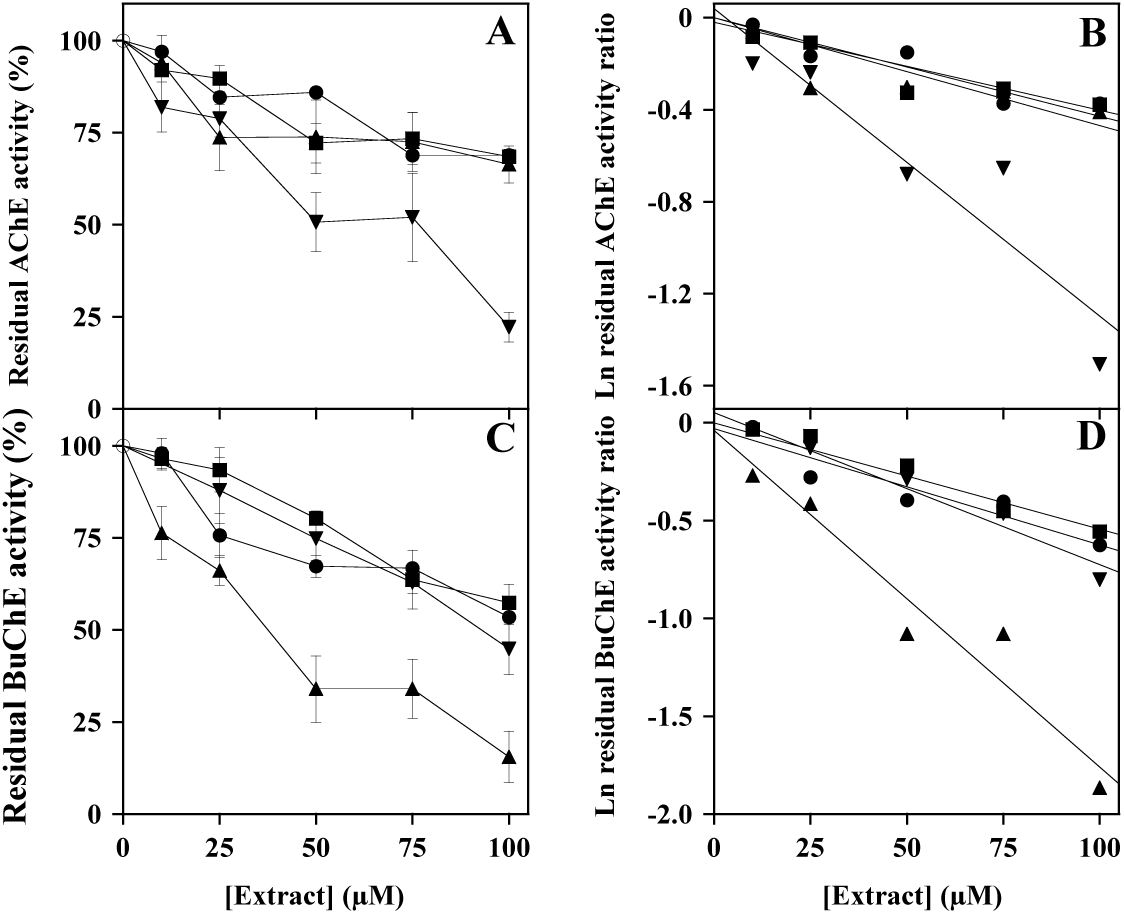
Effect of *Lo*CT, *Hc*CT, *Ms*F and *Ci*F extracts on the steady-state activity of AChE (**A**, **B**) and BuChE (**C**, **D**). The ratio of activity was assayed in the absence (μ) or in the presence of the indicated concentrations of *Lo*CT (λ), *Hc*CT (.&), *Ms*F (T) and *Ci*F (ν) and expressed as a percentage for AChE (**A**) and BuChE (**C**). The data obtained on four or three independent measurements for AChE or BuChE, respectively, were also analyzed after a logarithmic transformation of the activity ratio of AChE (**B**) and BuChE (**D**). The correlation coefficient *R* of the linear equation ranged between 0.909–0.964 (**B**) or 0.977–0.990 (**D**).

**Table 3.** Inhibition by polyphenolic extracts on the steady activity of cholinesterases.

| Extract | Acetylcholinesterase (AChE) |  | Butyrylcholinesterase (BuChE) |  |
| --- | --- | --- | --- | --- |
|  | Concentration interval | IC<sub>50</sub> | Concentration interval | IC<sub>50</sub> |
| | (μM) | (μM) | (μM) | (μM) |
| *Lo*CT | 0 – 100 | > 100 ($R = 0.964$) | 0 – 100 | > 100 ($R = 0.977$) |
| *Hc*CT | 0 – 100 | > 100 ($R = 0.909$) | 0 – 100 | $40 \pm 6$ ($R = 0.988$) |
| *Ms*F | 0 – 100 | $52 \pm 7$ ($R = 0.963$) | 0 – 100 | $89 \pm 8$ ($R = 0.990$) |
| *Ci*F | 0 – 100 | > 100 ($R = 0.960$) | 0 – 100 | > 100 ($R = 0.982$) |

The IC<sub>50</sub> values were extrapolated from a logarithmic transformation of the activity data obtained on four or three independent measurements for AChE or BuChE, respectively. The correlation coefficient $R$ of the straight lines used for calculation is reported in parentheses.

To get an insight in the inhibition mechanism of the extracts, kinetic measurements of the AChE activity were realized upon the addition of the various extracts. As shown in Fig. 4, their effect was evaluated using both a low, 20–40 µM, and a high, 100 µM, concentration of *Hc*CT, *Ms*F and *Ci*F; in the case of *Lo*CT, only the high concentration was used, because of the weak inhibition power by this extract. The *v*_i_ data of AChE were analyzed in the typical Michaelis–Menten representation (Figs. 4A, 4C, 4E, and 4G) and in Lineweaver–Burk plots (Figs. 4B, 4D, 4F, and 4H), thus allowing the extrapolation of the kinetic parameters *K_M_* and *V*_max_ in the absence or in the presence of the various plant extracts; the similar values obtained with both Michaelis–Menten and Lineweaver–Burk equations were averaged and reported in Table 4.

**Figure 4.**
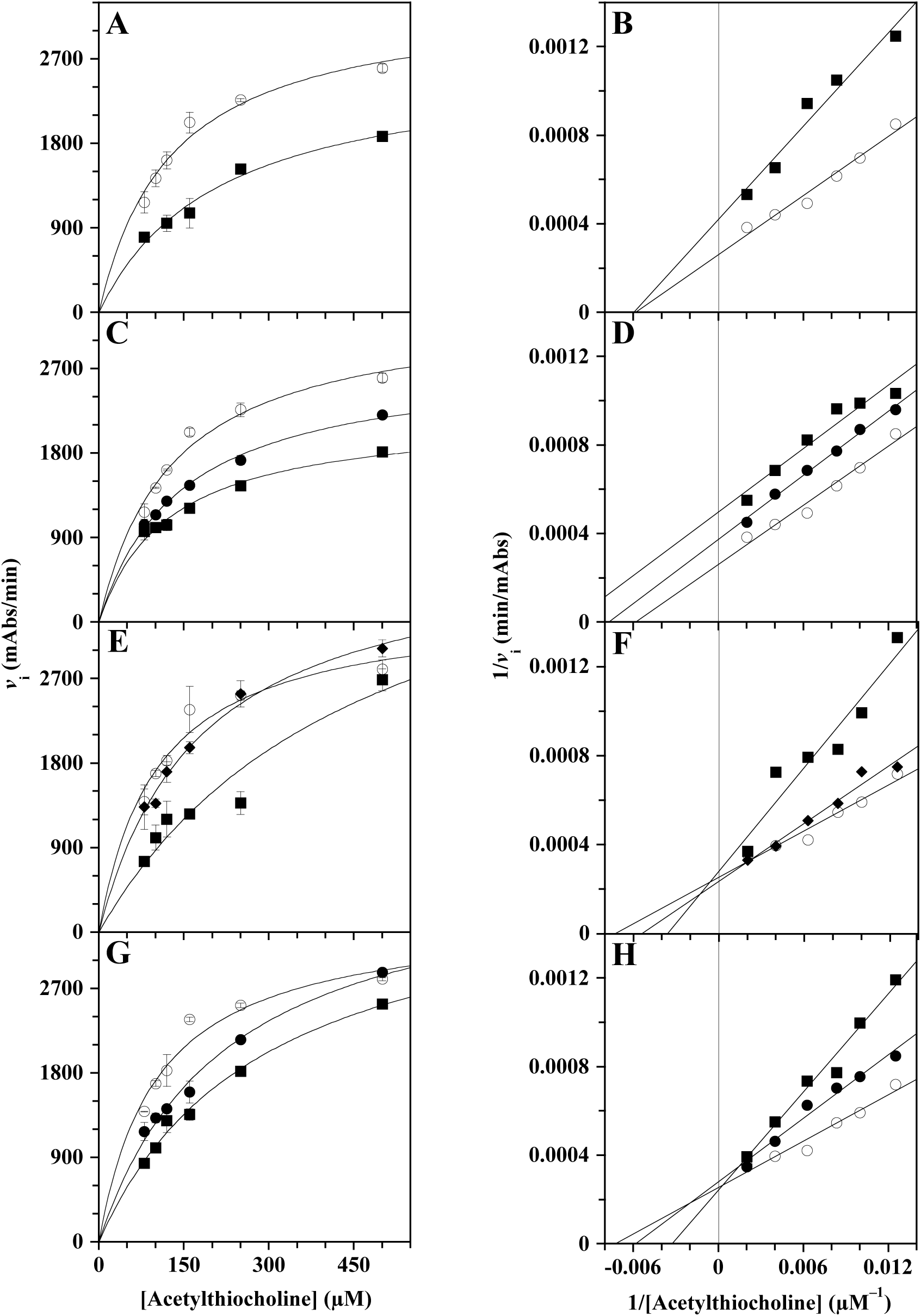
Kinetic analysis of the AChE inhibition by *Lo*CT, *Hc*CT, *Ms*F and *Ci*F extracts. The kinetic measurements of AChE activity were realized as reported in the Methods section in the presence of 80–500 µM acetylthiocholine concentration (three determinations for each concentration), without (μ) or with the following concentrations of polyphenolic extracts: (**A**, **B**) 100 µM (ν) *Lo*CT; (**C**, **D**) 20 µM (λ) or 100 µM (ν) *Hc*CT; (**E**, **F**) 40 µM (υ) or 100 µM (ν)) *Ms*F; (**G**, **H**) 20 µM (λ) or 100 µM (ν) *Ci*F. Data were reported reported as the initial velocity of substrate transformation (mean value ± S.E.) using the hyperbolic Michaelis–Menten equation (**A**, **C**, **E**, **G**) or the Lineweaver–Burk representation (**B**, **D**, **F**, **H**). The correlation coefficient *R* of the hyperbolic or linear equation ranged between 0.978–0.987 (**A**, **B**), 0.969–0.997 (**C**, **D**), 0.956–0.993 (**E**, **F**), 0.968– 0.994 (**G**, **H**).

**Table 4.** Effect of polyphenolic extracts on the kinetic parameters of AChE.

| Extract | Concentration<br>($\mu\text{M}$) | $K_M$ acetylthiocholine<br>($\mu\text{M}$)* | $V_{\max}$<br>(mAbs/min)* | Putative inhibition<br>mechanism | $K_i$<br>($\mu\text{M}$) | Calculation of<br>$K_i$ |
| --- | --- | --- | --- | --- | --- | --- |
| None | | $134 \pm 14$ | $3645 \pm 133$ (n = 8) | | | |
| LoCT | 100 | $167 \pm 22$ | $2528 \pm 154$ (n = 3) | non-competitive | $239 \pm 23$ | equation (b) |
| HcCT | 20 | $136 \pm 7$ | $2736 \pm 67$ (n = 4) | uncompetitive | $106 \pm 14$ | equations (c) |
| | 100 | $109 \pm 13$ | $2111 \pm 105$ (n = 4) | | | |
| MsF | 40 | $180 \pm 3$ | $4182 \pm 29$ (n = 3) | competitive | $83 \pm 12$ | equation (a) |
| | 100 | $373 \pm 99$ | $4265 \pm 733$ (n = 3) | | | |
| CiF | 20 | $197 \pm 27$ | $3832 \pm 282$ (n = 4) | competitive | $60 \pm 8$ | equation (a) |
| | 100 | $296 \pm 7$ | $4043 \pm 64$ (n = 4) | | | |

\*Number of replicates are reported in parentheses.

Indeed, *Lo*CT produced a decrease in the *V*_max_ of AChE activity with a minimum increase of the *K_M_*, whereas *Hc*CT caused a progressive decrease of the *V*_max_ concomitant to some decrease of the *K_M_*. On the other hand, *Ms*F and *Ci*F caused a progressive and consistent increase of the *K_M_*, without apparently affecting the *V*_max_ of the reaction. On the basis of these effects and using appropriate equations, we calculated the inhibition constant (*K*_i_) of each extract, an important parameter for measuring their inhibition power (Table 4). Indeed, *Ci*F (*K*_i_ = 60 ± 8 µM) had the greatest inhibition power, followed by *Ms*F (*K*_i_ = 83 ± 12 µM) and *Hc*CT (*K*_i_ = 106 ± 14 µM) in the order, and at a distance by *Lo*CT (*K*_i_ = 239 ± 23 µM). The inhibition mechanism displayed by the extracts was also evaluated, by inspecting the intersection of the straight lines obtained in Lineweaver–Burk plots. Indeed, the intersection on the abscissa axis with *Lo*CT (Fig. 4B) suggested a non-competitive inhibition mechanism, whereas the almost parallel lines obtained with *Hc*CT (Fig. 4D) pointed to an uncompetitive mechanism. Vice versa, the intersection on the ordinate axis with *Ms*F (Fig. 4F) and *Ci*F (Fig. 4H) indicated a competitive inhibition mechanism.

The same kinetic measurements were performed with the BuChE activity and the results are presented in Fig. 5. Also in this case, the *v*_i_ data obtained without or with the extracts was analyzed with the Michaelis–Menten (Figs. 5A, 5C, 5E, and 5G) and Lineweaver–Burk (Figs. 5B, 5D, 5F, and 5H) representation. The resulting values of *K_M_* and *V*_max_ reported in Table 5 suggest that *Lo*CT and *Ms*F provoked a decrease of the *V*_max_, without apparently affecting the *K_M_* of the reaction. On the other hand, *Hc*CT provoked a decrease of the *V*_max_ concomitant to an increase of the *K_M_*, whereas *Ci*F caused a decrease of both *V*_max_ and *K_M_* of BuChE activity. The corresponding values of *K*_i_ calculated for these extracts are reported in Table 5. Indeed, two extracts, namely *Hc*CT (*K*_i_ = 75 ± 13 µM) and *Ms*F (*K*_i_ = 97 ± 11 µM) were endowed with a moderate inhibition power towards BuChE, whereas *Lo*CT (*K*_i_ = 155 ± 6 µM) and *Ci*F (*K*_i_ = 141 ± 7 µM) showed a lower inhibition power. The inhibition mechanism was also evaluated through the Lineweaver–Burk plots. Indeed, the intersection of the straight lines on the abscissa axis with *Lo*CT (Fig. 5B) and *Ms*F (Fig. 5F) suggested their non-competitive inhibition mechanism. In the case of *Hc*CT (Fig. 5D), the lines intersected very close to the ordinate axis, thus suggesting a mixed mechanism, with some prevalence to a competitive inhibition. Finally, the almost parallel lines observed with *Ci*F (Fig. 5H) pointed to an uncompetitive mechanism.

**Figure 5.**
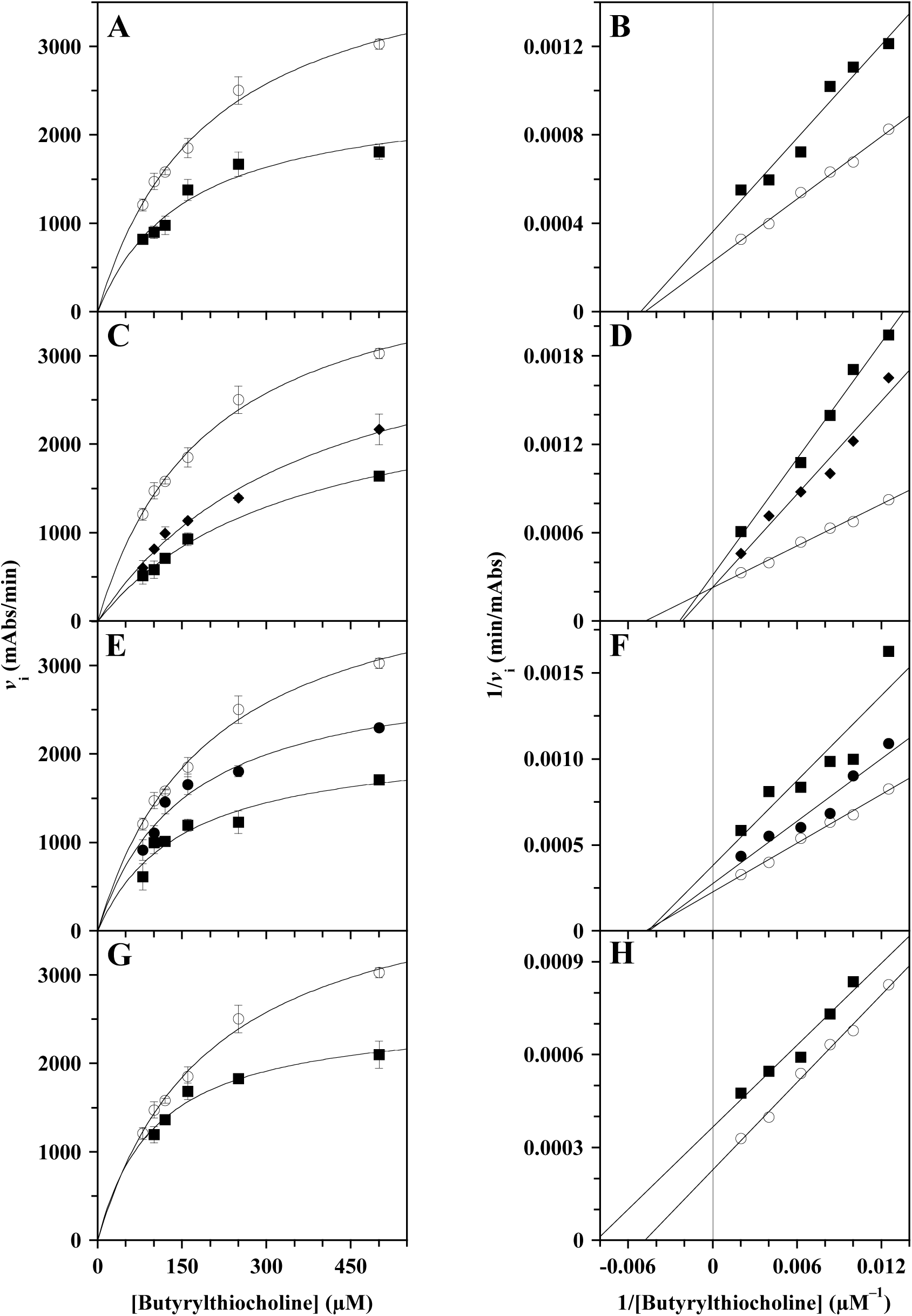
Kinetic analysis of the BuChE inhibition by *Lo*CT, *Hc*CT, *Ms*F and *Ci*F extracts. The kinetic measurements of BuChE activity were realized as reported in the Methods section in the presence of 80–500 µM butyrylthiocholine concentration (three determinations for each concentration), without (μ) or with the following concentrations of polyphenolic extracts: (**A**, **B**) 100 µM (ν) *Lo*CT; (**C**, **D**) 40 µM (υ) or 100 µM (ν) *Hc*CT; (**E**, **F**) 20 µM (λ) or 100 µM (ν)) *Ms*F; (**G**, **H**) 100 µM (ν) *Ci*F. Data were reported as the initial velocity of substrate transformation (mean value ± S.E.) using the hyperbolic Michaelis–Menten equation (**A**, **C**, **E**, **G**) or the Lineweaver–Burk representation (**B**, **D**, **F**, **H**). The correlation coefficient *R* of the hyperbolic or linear equation ranged between 0.976–0.996 (**A**, **B**), 0.982–0.997 (**C**, **D**), 0.907–0.996 (**E**, **F**), 0.977–0.996 (**G**, **H**).

**Table 5.** Effect of polyphenolic extracts on the kinetic parameters of BuChE.

| Extract | Concentration<br>($\mu\text{M}$) | $K_M$ butyrylthiocholine<br>($\mu\text{M}$)* | $V_{\max}$<br>(mAbs/min)* | Putative inhibition<br>mechanism | $K_i$<br>($\mu\text{M}$) | Calculation<br>of $K_i$ |
| --- | --- | --- | --- | --- | --- | --- |
| None | | $202 \pm 4$ | $4324 \pm 42$ (n = 4) | | | |
| LoCT | 100 | $175 \pm 19$ | $2623 \pm 133$ (n = 3) | non-competitive | $155 \pm 6$ | equation (b) |
| HcCT | 40 | $410 \pm 37$ | $3996 \pm 259$ (n = 3) | mixed, approaching<br>competitive | $75 \pm 13$ | equation (a) |
| | 100 | $380 \pm 33$ | $2968 \pm 173$ (n = 3) | | | |
| MsF | 20 | $186 \pm 32$ | $3314 \pm 294$ (n = 3) | non-competitive | $97 \pm 11$ | equation (b) |
| | 100 | $181 \pm 35$ | $2390 \pm 228$ (n = 3) | | | |
| CiF | 100 | $111 \pm 10$ | $2646 \pm 83$ (n = 3) | uncompetitive | $141 \pm 7$ | equations (c) |

\*Number of replicates are reported in parentheses.

### 3.3. Effect of the plant extracts on self-aggregation of the Aβ_1–40_ peptide and disaggregation of amyloid fibrils

To evaluate the effect of the extracts on amyloidogenesis of the Aβ_1–40_ peptide, we have analyzed if these samples might interfere with the Aβ_1–40_ self-aggregation process, as well as with the disaggregation of preformed amyloid fibrils (Fig. 6). Concerning the self-aggregation process, the Aβ_1–40_ peptide was incubated alone or in the presence of increasing concentration of the extracts and the extent of fibrils formation was reported in Fig. 6A. The data clearly show that all the four extracts interfered with the amyloid fibril formation process, with *Hc*CT exerting a slightly higher efficacy among the four extracts. After the logarithmic transformation of the amyloid fibril formation ratio, the resulting semi-logarithmic plots (Fig. 6B) allowed the extrapolation of the IC_50_ values for all extracts. This parameter confirmed that *Hc*CT (144 ± 6 µM) was the most effective in interfering fibrils formation, followed by *Ci*F and *Lo*CT (205 ± 12 µM and 207 ± 10 µM, respectively), and by *Ms*F (364 ± 10 µM) in the order (Table 6).

**Figure 6.**
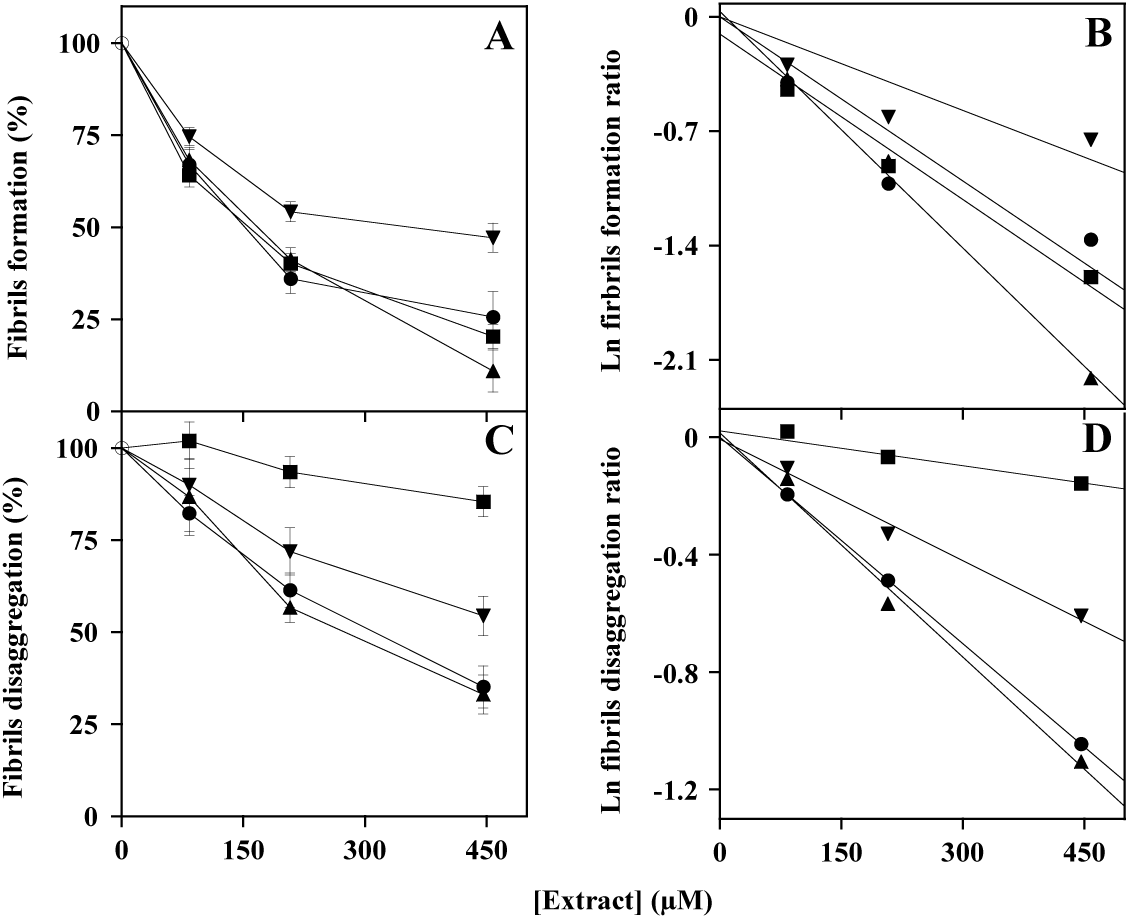
Effect of *Lo*CT, *Hc*CT, *Ms*F and *Ci*F extracts on amyloid fibrils formation (**A**, **B**) or disaggregation (**C**, **D**) process. The ratio of formation/disaggregation was measured in the absence (μ) or in the presence of the indicated concentrations of *Lo*CT (λ), *Hc*CT (.&), *Ms*F (T) and *Ci*F (ν) and expressed as a percentage for formation (**A**) or disaggregation (**C**). The data of three independent measurements for fibril formation or disaggregation, respectively, were also analyzed after a logarithmic transformation of the formation ratio (**B**) or disaggregation ratio (**D**). The correlation coefficient *R* of the linear equation ranged between 0.936–0.999 (**B**) or 0.980–1.000 (**D**).

**Table 6.**
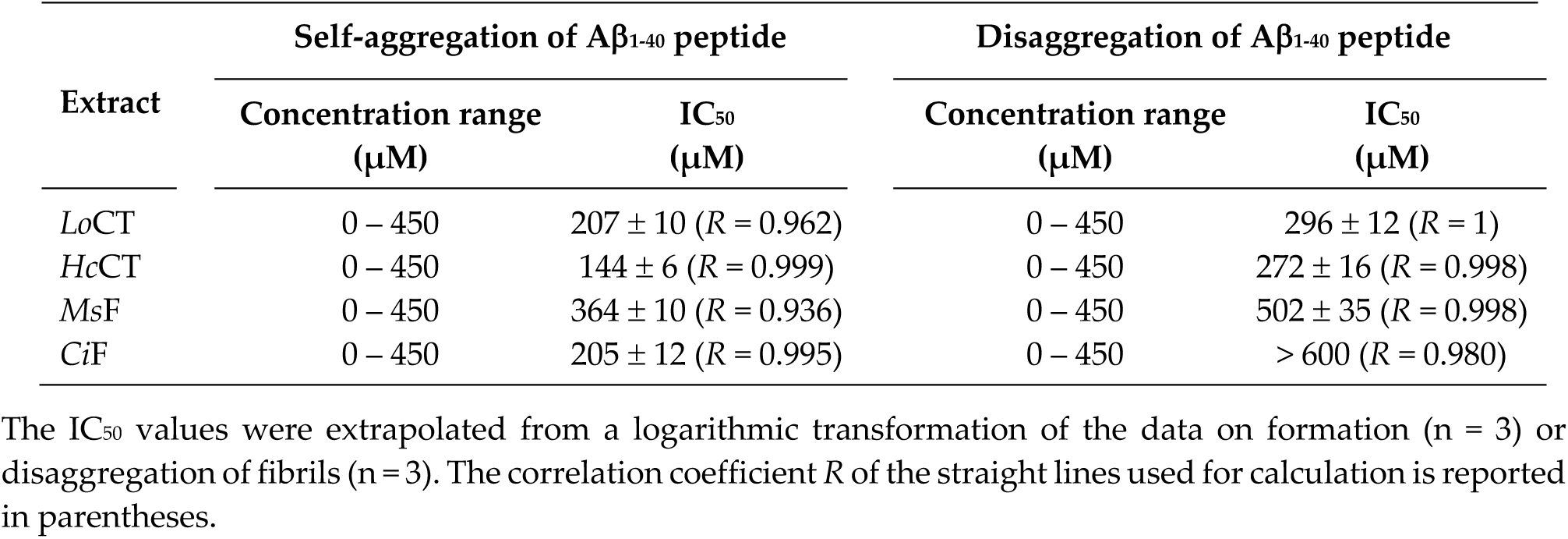
Effect of polyphenolic extracts on the amyloid fibril aggregation and disaggregation process.

Then, we moved to evaluate the effects exerted by the extracts on the amyloid disaggregation process (Fig. 6C). In this assay, all the extracts displayed a lower interfering effect compared to that observed in fibrils formation, although with a different order of effectiveness among them. To better address this point, the data were also analysed through the semi-logarithmic plots (Fig. 6D), thus allowing the extrapolation of the IC_50_ values reported in Table 6. Indeed, in the fibril disaggregation process, the lowest IC_50_ was found with *Hc*CT (272 ± 16 µM), closely followed by *Lo*CT (296 ± 12 µM), then by *Ms*F (502 ± 35 µM), and at a distance by *Ci*F (> 600 µM).

### 3.4. Effect of the plant extracts on the viability of a neuroblastoma cell line

The human neuroblastoma SH-SY5Y cell line was chosen to evaluate the effect of the polyphenolic extracts on cell viability of these cancer cells. To this aim, SH-SY5Y were exposed, for 24-h treatment, to increasing concentrations of the various extracts up to 250 µM for *Lo*CT, *Hc*CT and *Ms*F or up to 150 µM for *Ci*F. As vehicle, the DMSO concentration, was kept always lower than 0.6% (v/v). Cell viability was evaluated with the 3-(4,5-dimethylthiazol-2-yl)-2,5-biphenyltetrazolium bromide (MTT) assay and the results were reported in Fig. 7. The greatest effect was observed with *Ms*F, with an evident dose-dependent reduction of cell viability (Fig. 7C). In particular, after treatment with 250 µM *Ms*F, the cell viability of SH-SY5Y was reduced to 29%. The other extracts were much less effective and none of them reached 50% reduction of cell viability, even with the maximum dose of treatment. In particular, cell viability of SH-SY5Y was reduced to 74%, 66%, or 77%, after treatment with 250 µM *Lo*CT (Fig. 7A), 250 µM *Hc*CT (Fig. 7B), or 150 µM *Ci*F (Fig. 7D), respectively.

**Figure 7.**
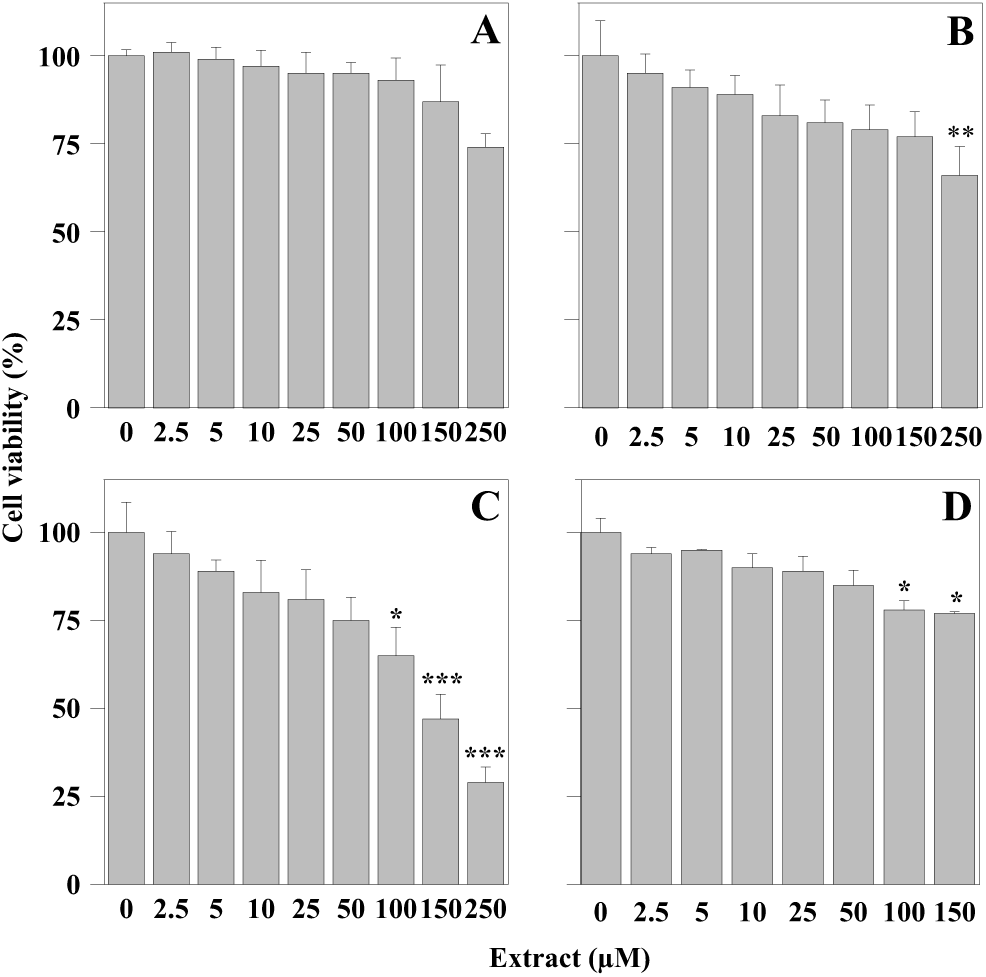
Cell viability of human neuroblastoma SH-SY5Ycell line after treatment with polyphenolic extracts. Cells were treated for 24 h with the indicated concentrations of *Lo*CT (panel **A**), *Hc*CT (panel **B**), *Ms*F (panel **C**), or *Ci*F (panel **D**). Control cells were incubated with 0.6 % (v/v) DMSO as a vehicle. Cell viability was determined with the MTT assay, as reported in Materials and Methods. The values, reported as a percentage compared to control cells, represent the mean ± standard error of separate experiments performed in triplicates. The significance was evaluated with *p* < 0.05 (*), 0.01 (**), and 0.001 (***).

## 4. DISCUSSION

In the last decades, natural polyphenols contained in edible plants have attracted the attention of many researchers for their possible benefits as adjutants of therapies against several human pathologies [36]. Indeed, it is known that an enriched-polyphenol diet is recommended to prevent and alleviate symptoms of neurodegenerative disorders, like AD and PD [37–40]. Polyphenols include a wide variety of different structures, comprising phenolic acids, flavonoids, lignans, and stilbenes; this variety highlights their complex biological functions. Understanding the origin and structural diversity of polyphenolic compounds is crucial for recognizing their antioxidant, anti-inflammatory and neuroprotective properties.

In a recent paper, we demonstrated that polyphenols extracted from Mediterranean forage crops, such as *L. ornithopodioides*, *H. coronarium*, *M. sativa*, and *C. intybus*, display antioxidant activities, hopefully useful for the design of drugs beneficial for human health [32]. In the present work, composition of these extracts, namely *Lo*CT, *Hc*CT, *Ms*F and *Ci*F, was analysed in detail and the different components, including 24 phenolic acids and 25 flavonoids, were overall identified. The identification of natural compounds in extraction mixtures constitutes an indispensable information for a future analysis and focus on the effects by single components on various metabolic pathways. However at moment, the working hypothesis followed in the present work is that the effects observed by plant extracts could be due to a specific combination of their various components. Indeed, several dietary regimes, including the popular Mediterranean diet, place at the top the usage of plants as natural foods and/or coadjutants of therapeutic treatments, without considering the effects of each specific component [16,18–21].

Because of the neuroprotective properties possessed by polyphenols [22–24], we decided to investigate on the effects caused by *Lo*CT, *Hc*CT, *Ms*F and *Ci*F on two crucial enzymes of the neurotransmission process, such as AChE and BuChE. Indeed, all extracts acted as moderate inhibitors of both enzymes with *K*_i_ values ranging in the 60 – 240 µM interval. Among the four samples, the flavonoid-containing extracts had a slightly greater potency towards both cholinesterases, compared to samples containing condensed tannins. *Ms*F had an almost similar inhibition strength against AChE (*K*_i_ = 83 µM) and BuChE (*K*_i_ = 97 µM), whereas *Ci*F displayed a greater inhibition towards AChE (*K*_i_ = 60 µM) compared to BuChE (*K*_i_ = 141 µM). On the other hand, despite of their lower strength, extracts with condensed tannins showed a preference for BuChE inhibition, with *K*_i_ values of 75 µM and 155 µM for *Hc*CT and *Lo*CT, respectively. The study of the effects on kinetic parameters of both cholinesterase activities revealed differences in the inhibition mechanism exerted by extracts, that are of difficult interpretation. For instance, *Ms*F and *Ci*F had the same competitive mechanism towards AChE, whereas they showed different mechanisms towards BuChE, *i.e.* non-competitive and uncompetitive for *Ms*F and *Ci*F, respectively. This different behavior could be explained with the combination/overlapping of different inhibition mechanisms exhibited by single components in the extracts.

Another parameter considered for evaluating the possible effects by plant extracts on neurodegenerative diseases was the formation and disassembly of amyloid fibrils deriving from the aggregation of Aβ_1-40_ peptide. Our data on self-aggregation process indicate that all extracts acted as moderate inhibitors of this process, with IC_50_ values ranging in a moderately wide interval, 144 – 364 µM; indeed, the greatest efficacy found with *Hc*CT was not so distant from that observed with *Ms*F. Concerning the effects on disaggregation of amyloid fibrils, all extracts displayed a common lower efficacy compared to the self-aggregation process. Furthermore, greater differences were found in the interval of IC_50_ values, ranging from 272 µM to > 600 µM. However, among the various extracts, the *Hc*CT sample had the greatest efficacy in both Aβ_1–40_ peptide self-aggregation and disaggregation process. All these results are congruent with recent studies reporting that plant derived extracts/compounds can protect neuronal cell damage using an *in vitro* model of AD, by interfering with the Aβ_1–40_ peptide aggregation [11, 41–43], although the Authors underlined the hindering aspects related to the intracellular bioavailabilty and the capability to cross the ematoencephalic barrier by substances of natural origin.

We have also checked the cytotoxicity against human neuroblastoma cells, as the usage of polyphenols is probably associated to a reduced incidence of several human pathologies [44,45]. Indeed, we have already reported that the extract from *M. sativa* affected the cell viability of the gastric MKN-28 and AGS cancer lines [32]. Under this regard, the neuroblastoma cell line SH-SY5Y, mimicking immature cholinergic neurons [11,46], is frequently used as an *in vitro* model of AD. We have found that *Ms*F significantly reduced the viability of SH-SY5Y cells, a finding potentially useful for considering natural polyphenolic extracts in cancer therapy. However, comparing the cytotoxicity against neuroblastoma cells with the IC_50_ values for enzyme inhibition and amyloidogenesis, the therapeutic window is restricted to a narrow interval, thus limiting a practical application of these Mediterranean forage crops extracts.

## 5. CONCLUSION

The results reported in this study indicate that, although to a different extent, *L. ornithopodioides*, *H. coronarium, M. sativa*, and *C. intybus* leaf extracts, inhibit AChE and BuChE, and interfere with the peptide Aβ_1–40_ amyloidogenesis process; therefore, they have properties for postulating the usage as adjutants in the treatment of AD. On the other hand, the reduced cell viability caused by *M. sativa* extract in a neuroblastoma cell line, as well as in gastric cancer cell lines [32], suggests its potential use as a coadjutant in anticancer therapies; however, further investigation is required to assess this hypothesis. In particular, the intracellular bioavailability of the extract components, and their ability to cross the ematoencephalic barrier should be investigated, to establish the potential use of these substances in therapies. All these findings strengthen the consideration that polyphenols can serve as models for developing innovative strategies aimed at the prevention of human diseases and enhancing human health [18,22].

## Supporting information

Supplementary_D_Errico_et_al_2025

## REFERENCES

1. Zanetti O, Solerte SB, Cantoni F. Life expectancy in Alzheimer’s disease (AD). Arch Gerontol Geriatr. 2009; 49 Suppl. 1: 237–243. Doi: 10.1016/j.archger.2009.09.035

2. Golbe LI, Leyton CE. Life expectancy in Parkinson disease. Neurology. 2018; 91: 991–992. Doi: 10.1212/WNL.0000000000006560

3. Caligiore D, Giocondo F, Silvetti M. The neurodegenerative elderly syndrome (NES) hypothesis: Alzheimer and Parkinson are two faces of the same disease. IBRO Neurosci Rep. 2022; 13: 330–343. Doi: 10.1016/j.ibneur.2022.09.007

4. Chen JJ. Parkinson’s disease: health-related quality of life, economic cost, and implications of early treatment. Am J Manag Care. 2010; 16 Suppl. Implications: S87–93.

5. del Campo M, Peeters CFW, Johnson ECB, Vermunt L, Hok-A-Hin YS, van Nee M et al. CSF proteome profiling across the Alzheimer’s disease spectrum reflects the multifactorial nature of the disease and identifies specific biomarker panels. Nat Aging. 2022; 2: 1040–1053. Doi: 10.1038/s43587-022-00300-1

6. Gątarek P, Kałużna-Czaplińska J. Integrated metabolomics and proteomics analysis of plasma lipid metabolism in Parkinson’s disease. Expert Rev Proteomics. 2024; 21: 13–25. Doi: 10.1080/14789450.2024.2315193

7. Van Bulck M, Sierra-Magro A, Alarcon-Gil J, Perez-Castillo A, Morales-Garcia JA. Novel approaches for the treatment of Alzheimer’s and Parkinson’s disease. Int J Mol Sci. 2019; 20: 719. Doi: 10.3390/ijms20030719

8. Lane RM, Potkin SG, Enz A. Targeting acetylcholinesterase and butyrylcholinesterase in dementia. Int J Neuropsychopharmacol. 2006; 9: 101–124. Doi: 10.1017/S1461145705005833

9. Schliebs R, Arendt T. The cholinergic system in aging and neuronal degeneration. Behav Brain Res. 2011; 221: 555–563. Doi: 10.1016/j.bbr.2010.11.058

10. Ballard CG, Greig NH, Guillozet-Bongaarts AL, Enz A, Darvesh S. Cholinesterases: roles in the brain during health and disease. Curr Alzheimer Res. 2005; 2: 307–318. Doi: 10.2174/1567205054367838

11. Maiuolo J, Costanzo P, Masullo M, D’Errico A, Nasso R, Bonacci S et al. Hydroxytyrosol-donepezil hybrids play a protective role in an *in vitro* induced Alzheimer’s disease model and in neuronal differentiated human SH-SY5Y neuroblastoma cells. Int J Mol Sci. 2023; 24: 13461. Doi: 10.3390/ijms241713461

12. d’Angremont E, Begemann MJH, van Laar T, Sommer IEC. Cholinesterase inhibitors for treatment of psychotic symptoms in Alzheimer disease and Parkinson disease: a meta-analysis. JAMA Neurol. 2023; 80: 813–823. Doi: 10.1001/jamaneurol.2023.1835

13. Azargoonjahromi A. The duality of amyloid-β: its role in normal and Alzheimer’s disease states. Mol Brain. 2024; 17: 44. Doi: 10.1186/s13041-024-01118-1

14. Pagano K, Tomaselli S, Molinari H, Ragona L. Natural compounds as inhibitors of Aβ peptide aggregation: chemical requirements and molecular mechanisms. Front Neurosci. 2020; 14: 619667. Doi: 10.3389/fnins.2020.619667

15. Hansen RA, Gartlehner G, Webb AP, Morgan LC, Moore CG, Jonas DE. Efficacy and safety of donepezil, galantamine, and rivastigmine for the treatment of Alzheimer’s disease: a systematic review and meta-analysis. Clin Interv Aging. 2008; 3: 211–225

16. Kumar A, Tiwari A, Sharma A. Changing paradigm from one target one ligand towards multi-target directed ligand design for key drug targets of Alzheimer disease: an important role of in silico methods in multi-target directed ligands design. Curr Neuropharmacol. 2018; 16: 726– 739. Doi: 10.2174/1570159X16666180315141643

17. Campos-Esparza MR, Torres-Ramos MA. Neuroprotection by natural polyphenols: molecular mechanisms. Cent Nerv Syst Agents Med Chem. 2010; 10: 269–277. Doi: 10.2174/187152410793429728

18. Nasso R, Pagliara V, D’Angelo S, Rullo R, Masullo M, Arcone R. *Annurca* apple polyphenol extract affects acetyl-cholinesterase and mono-amine oxidase in vitro enzyme activity. Pharmaceuticals (Basel). 2021; 14: 62. Doi: 10.3390/ph14010062

19. Silva RFM, Pogačnik L. Polyphenols from food and natural products: neuroprotection and safety. Antioxidants (Basel). 2020; 9: 61. Doi: 10.3390/antiox9010061

20. Arias-Sánchez RA, Torner L, Fenton Navarro B. Polyphenols and neurodegenerative diseases: potential effects and mechanisms of neuroprotection. Molecules. 2023; 28: 5415. Doi: 10.3390/molecules28145415

21. Ullah A, Munir S, Badshah SL, Khan N, Ghani L, Poulson BG et al. Important flavonoids and their role as a therapeutic agent. Molecules. 2020; 25: 5243. Doi: 10.3390/molecules25225243

22. Arcone R, D’Errico A, Nasso R, Rullo R, Poli A, Di Donato P et al. Inhibition of enzymes involved in neurodegenerative disorders and Aβ_1-40_ aggregation by *Citrus limon* peel polyphenol extract. Molecules. 2023; 28: 6332. Doi: 10.3390/molecules28176332

23. Orhan I, Aslan S, Kartal M, Şener B, Hüsnü Can Başer K. Inhibitory effect of Turkish Rosmarinus officinalis L. on acetylcholinesterase and butyrylcholinesterase enzymes. Food Chem. 2008; 108: 663–668. Doi: 10.1016/j.foodchem.2007.11.023

24. de Torre MP, Cavero RY, Calvo MI. Anticholinesterase activity of selected medicinal plants from Navarra region of Spain and a detailed phytochemical investigation of *Origanum vulgare* L. ssp. *vulgare*. Molecules. 2022; 27: 7100. Doi: 10.3390/molecules27207100

25. Tava A, Biazzi E, Ronga D, Pecetti L, Avato P. Biologically active compounds from forage plants. Phytochem Rev. 2022; 21: 471–501. Doi.org/10.1007/s11101-021-09779-9

26. Usman M, Khan WR, Yousaf N, Akram S, Murtaza G, Kudus KA et al. Exploring the hytochemicals and anti-cancer potential of the members of *Fabaceae* family: a comprehensive review. Molecules. 2022; 27: 3863. Doi: 10.3390/molecules27123863

27. Burlando B, Pastorino G, Salis A, Damonte G, Clericuzio M, Cornara L. The bioactivity of *Hedysarum coronarium* extracts on skin enzymes and cells correlates with phenolic content. Pharm Biol. 2017; 55: 1984–1991. Doi: 10.1080/13880209.2017.1346691

28. Bora KS, Sharma A. Evaluation of antioxidant and cerebroprotective effect of *Medicago sativa* Linn. against ischemia and reperfusion insult. Evid Based Complement Alternat Med. 2011; 792167. Doi: 10.1093/ecam/neq019

29. Raeeszadeh M, Mortazavi P, Atashin-Sadafi R. The ntioxidant, anti-inflammatory, pathological, and behavioural effects of *Medicago sativa L.* (Alfalfa) extract on brain injury caused by nicotine in male rats. Evid Based Complement Alternat Med. 2021; 6694629. Doi: 10.1155/2021/6694629

30. Rolnik A, Olas B. The plants of the *Asteraceae* family as agents in the protection of human health. Int J Mol Sci. 2021; 22: 3009. Doi: 10.3390/ijms22063009

31. Perović J, Tumbas Šaponjac V, Kojić J, Krulj J, Moreno DA, García-Viguera C et al. Chicory (*Cichorium intybus* L.) as a food ingredient - Nutritional composition, bioactivity, safety, and health claims: A review. Food Chem. 2021; 336: 127676. Doi: 10.1016/j.foodchem.2020.127676

32. Rullo R, Nasso R, D’Errico A, Biazzi E, Tava A, Landi N et al. Effect of polyphenolic extracts from leaves of Mediterranean forage crops of enzymrs involved in the oxidative stress, and useful for alternative cancer treatments. J Appl Pharm Sci. 2025; 15: in press

33. Scacco S, Acquaviva S, França Vieira e Silva F, Zhang JH, Lo Muzio L, Corso G et al. Bioactivity and Neuroprotective Effects of Extra Virgin Olive Oil in a Mouse Model of Cerebral Ischemia: An In Vitro and In Vivo Study. Int J Mol Sci. 2025; 26, 1771–1802.

34. Vitale RM, Morace AM, D’Errico A, Ricciardi F, Fusco A, Boccella S et al. Identification of cannabidiolic and cannabigerolic acids as MTDL AChE, BuChE, and BACE-1 inhibitors against Alzheimer’s disease by in silico, in vitro, and in vivo studies. Phytother Res. 2025; 39: 233–245. Doi: 10.1002/ptr.8369

35. Rullo R, Cerchia C, Nasso R, Romanelli V, De Vendittis E, Masullo M et al. Novel reversible inhibitors of xanthine oxidase targeting the active site of the enzyme. Antioxidants (Basel). 2023; 12: 825. Doi: 10.3390/antiox12040825

36. Jalouli M, Rahman MA, Biswas P, Rahman H, Harrath AH, Lee I-S et al. Targeting natural antioxidant polyphenols to protect neuroinflammation and neurodegenerative diseases: a comprehensive review. Front Pharmacol. 2025; 16: 1492517. Doi: 10.3389/fphar.2025.1492517

37. Carregosa D, Mota S, Ferreira S, Alves-Dias B, Loncarevic-Vasiljkovic N, Crespo CL et al. Overview of beneficial effects of (poly)phenol metabolites in the context of neurodegenerative diseases on model organisms. Nutrients. 2021; 13: 2940. Doi: 10.3390/nu13092940

38. Spencer JPE. Food for thought: The role of dietary flavonoids in enhancing human memory, learning and neuro-cognitive performance. Proc Nutr Soc. 2008; 67: 238–252. Doi: 10.1017/S0029665108007088

39. Psaltopoulou T, Sergentanis TN, Panagiotakos DB, Sergentanis IN, Kosti R, Scarmeas N. Mediterranean diet, stroke, cognitive impairment and depression: A meta-analysis. Ann Neurol. 2013; 74: 580–591. Doi: 10.1002/ana.23944

40. Mayr HL, Thomas CJ, Tierney AC, Kucianski T, George ES, Ruiz-Canela M et al. Randomization to 6-month Mediterranean diet compared with a low-fat diet leads to improvement in the dietary inflammatory index scores in patients with coronary heart disease: The AUSMED heart trial. Nutr Res. 2018; 55: 94–107. Doi: 10.1016/j.nutres.2018.04.006

41. Pagano K, Tomaselli S, Molinari H, Ragona L. Natural compounds as inhibitors of Aβ peptide aggregation: chemical requirements and molecular mechanisms. Front Neurosci. 2020; 14: 619667. Doi: 10.3389/fnins.2020.619667

42. Li F, Zhan C, Dong X, Wei G. Molecular mechanisms of resveratrol and EGCG in the inhibition of Aβ_42_ aggregation and disruption of Aβ_42_ protofibril: similarities and differences. Phys Chem Chem Phys. 2021; 23: 18843–18854. Doi: 10.1039/d1cp01913a

43. Sun Y, Wang X, Zhang X, Li Y, Wang D, Sun F et al. Di-caffeoylquinic acid: a potential inhibitor for amyloid-beta aggregation. J Nat Med. 2024; 78: 1029–1043. Doi: 10.1007/s11418-024-01825-y

44. El-Saadony MT, Yang T, Saad AM, Alkafaas SS, Elkafas SS, Eldeeb GS et al. Polyphenols: Chemistry, bioavailability, bioactivity, nutritional aspects and human health benefits: A review. Int J Biol Macromol. 2024; 277: 134223. Doi: 10.1016/j.ijbiomac.2024.134223

45. Sun S, Liu Z, Lin M, Gao N, Wang X. Polyphenols in health and food processing: antibacterial, anti-inflammatory, and antioxidant insights. Front Nutr. 2024; 11: 1456730. Doi: 10.3389/fnut.2024.1456730

46. Dubey SK, Ram MS, Krishna KV, Saha RN, Singhvi G, Agrawal M et al. Recent expansions on cellular models to uncover the scientific barriers towards drug development for Alzheimer’s disease. Cell Mol Neurobiol. 2019; 39: 181–209. Doi: 10.1007/s10571-019-00653-z

