## Supplementary_D_Errico_et_al_2025 for "Effects of Polyphenolic Extracts from Mediterranean Forage Crops on Cholinesterases and Amyloid Aggregation Relevant to Neurodegenerative Diseases"

**Supplementary Figure S1**. Extract Ion Chromatogram (XIC) of Multi Reaction Monitoring (MRM) of the optimized 43 Phenolic acids.

**Supplementary Figure S2**. Extract Ion Chromatogram (XIC) of Multi Reaction Monitoring (MRM) of the optimized 38 Flavonoids.

**Supplementary Table S1.** Optimized Q1 mass, product ion and parameters for sMRM experiment of Phenolic acids and Flavonoids.

**Supplementary Table S2**. Linearity and sensitivity data for phenolic acids and flavonoids.

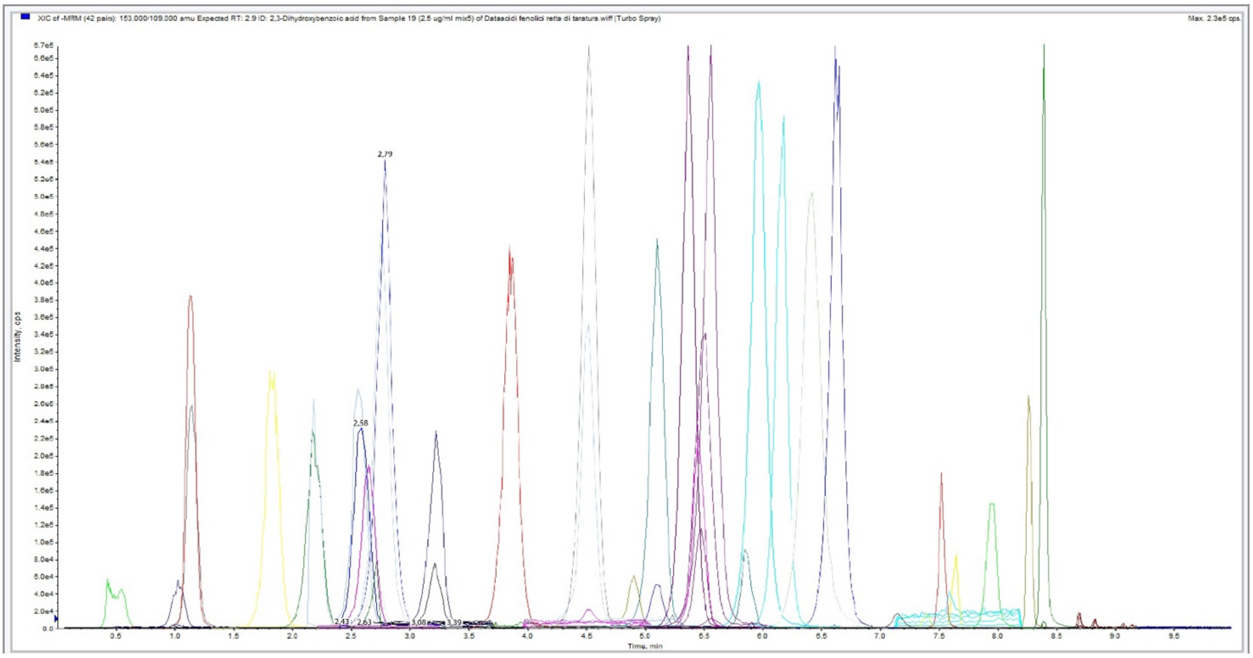


**Supplementary Figure S1**. Extract Ion Chromatogram (XIC) of Multi Reaction Monitoring (MRM) of the optimized 43 Phenolic acids.

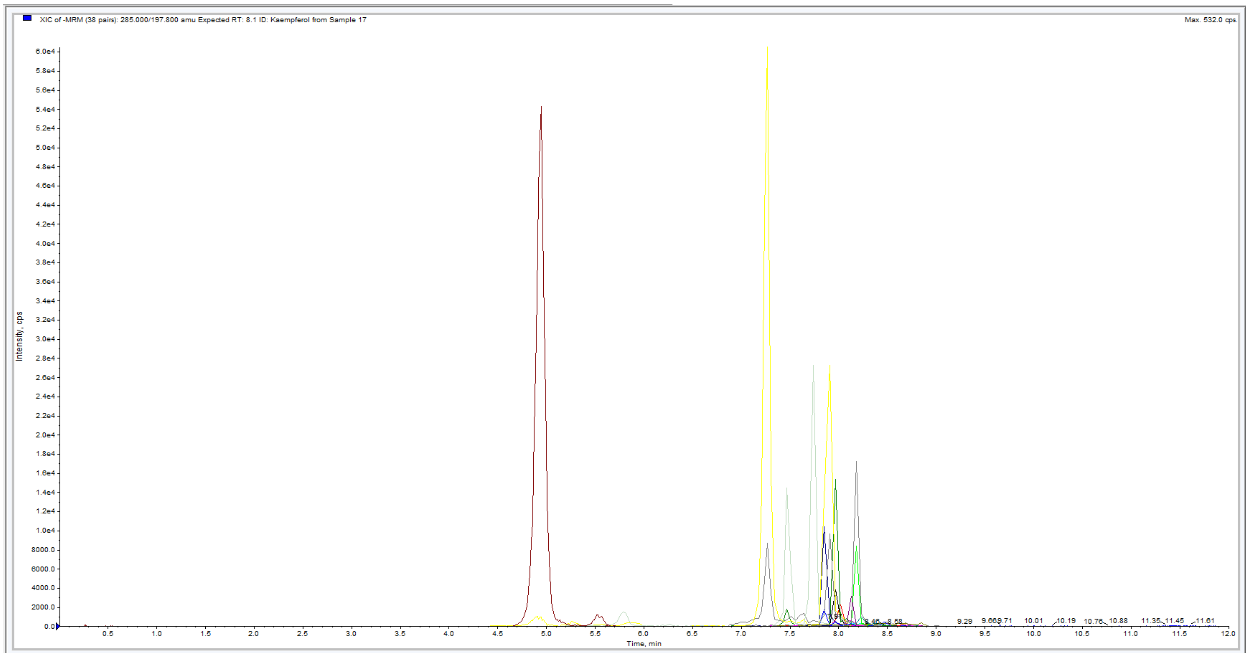


**Supplementary Figure S2**. Extract Ion Chromatogram (XIC) of Multi Reaction Monitoring (MRM) of the optimized 38 Flavonoids.

**Supplementary Table S1.** Optimized Q1 mass, product ion and parameters for sMRM experiment of Phenolic acids and Flavonoids.

| Metabolite | Metabolite Class[a] | Precursor ion (*m/z*) | Product ion (*m/z*) | DP[b] | EP[c] | CE[d] | CXP[e] | RT[f] (min) |
|---|---|---|---|---|---|---|---|---|
| Quinaldic Acid | IS | 174.05 $[M+H]^+$ | 128.0 | 70 | 10 | 30 | 9 | 2.57 |
| o-Anisic Acid | PA | 152.8 $[M+H]^+$ | 135.1 | 70 | 10 | 17 | 20 | 4.78 |
| 2,3-Dihydroxybenzoic acid | PA | 153.0 $[M-H]^-$ | 109.0 | -49 | -10.9 | -19 | -5 | 2.94 |
| Chlorogenic acid | PA | 353.1 $[M-H]^-$ | 190.8 | -49 | -10.9 | -20 | -10 | 3.99 |
| Syringic acid | PA | 197.0 $[M-H]^-$ | 152.9 | -49 | -10.9 | -16 | -16 | 4.17 |
| p-Coumaric acid | PA | 163.0 $[M-H]^-$ | 119.1 | -49 | -10.9 | -12 | -10 | 4.72 |
| m-Hydrocoumaric acid | PA | 165.0 $[M-H]^-$ | 120.6 | -49 | -10.9 | -18 | -7 | 4.78 |
| Ferulic acid | PA | 193.1 $[M-H]^-$ | 134.1 | -49 | -10.9 | -20 | -4 | 5.62 |
| Sinapic acid | PA | 223.1 $[M-H]^-$ | 163.5 | -49 | -10.9 | -19 | -16 | 6.01 |
| Aspirin | PA | 179.1 $[M-H]^-$ | 136.7 | -49 | -10.9 | -12 | -9 | 5.72 |
| *trans*-Cinnamic acid | PA | 147.0 $[M-H]^-$ | 103.0 | -49 | -10.9 | -14 | -10 | 7.66 |
| 4-Hydroxybenzoic acid | PA | 137.0 $[M-H]^-$ | 92.7 | -49 | -10.9 | -18 | -7 | 1.84 |
| 2,6-Dihydroxybenzoic acid | PA | 153.0 $[M-H]^-$ | 108.7 | -49 | -10.9 | -18 | -19 | 6.44 |
| Dihydrocaffeic acid | PA | 181.1 $[M-H]^-$ | 162.8 | -49 | -10.9 | -19 | -15 | 7.91 |
| Caffeic acid | PA | 179.0 $[M-H]^-$ | 134.4 | -49 | -10.9 | -20 | -7.5 | 3.17 |
| Phloretic acid | PA | 165.1 $[M-H]^-$ | 148.7 | -49 | -10.9 | -20 | -9 | 8.67 |
| Hydroferulic acid | PA | 195.1 $[M-H]^-$ | 135.7 | -49 | -10.9 | -15 | -5.6 | 4.9 |
| Ellagic acid dihydrate | PA | 301.0 $[M-H]^-$ | 144.8 | -49 | -10.9 | -43 | -17 | 7.1 |
| 5-Methoxysalicylic acid | PA | 167.0 $[M-H]^-$ | 107.6 | -49 | -10.9 | -30 | -14 | 6.6 |
| Catechol | PA | 109.0 $[M-H]^-$ | 108.9 | -49 | -10.9 | -11 | -31 | 1.1 |
| Gentisic acid | PA | 153.0 $[M-H]^-$ | 107.7 | -49 | -10.9 | -27 | -7 | 2.19 |
| 4-Acetocatechol | PA | 151.0 $[M-H]^-$ | 107.3 | -49 | -10.9 | -27 | -9 | 2.64 |
| 4-Methylcatechol | PA | 123.0 $[M-H]^-$ | 94.5 | -49 | -10.9 | -22 | -14 | 2.9 |
| 2,6-Dimethoxybenzoic acid | PA | 181.1 $[M-H]^-$ | 136.7 | -49 | -10.9 | -10.6 | -9 | 4.5 |
| Acetylphloroglucinol | PA | 167.0 $[M-H]^-$ | 122.8 | -49 | -10.9 | -23 | -11 | 5.05 |
| Salicylic acid | PA | 137.0 $[M-H]^-$ | 92.6 | -49 | -10.9 | -25 | -7 | 5.54 |
| trans-2-Hydroxycinnamic acid | PA | 163.0 $[M-H]^-$ | 119.2 | -49 | -10.9 | -15 | -26 | 6.14 |
| Caffeic acid dimethyl ether | PA | 207.0 $[M-H]^-$ | 102.7 | -49 | -10.9 | -19 | -16 | 7.34 |
| 3-Methoxyhydrocinnamic acid | PA | 179.1 $[M-H]^-$ | 119.9 | -49 | -10.9 | -17.5 | -13 | 7.48 |
| Gallic acid | PA | 169.0 $[M-H]^-$ | 124.9 | -49 | -10.9 | -21 | -9 | 0.52 |
| 3,5-Dihydroxybenzoic acid | PA | 153.0 $[M-H]^-$ | 108.8 | -49 | -10.9 | -15 | -9 | 1.02 |
| Vanillic acid | PA | 167.0 $[M-H]^-$ | 151.8 | -49 | -10.9 | -18 | -14 | 3.07 |
| Nordihydroguaiaretic Acid | PA | 301.1 $[M-H]^-$ | 121.6 | -49 | -10.9 | -34.5 | -23 | 8.25 |
| Terephthalic acid | PA | 165.0 $[M-H]^-$ | 120.5 | -49 | -10.9 | -14 | -12 | 3.19 |
| 4-Acetylresorcinol | PA | 151.0 $[M-H]^-$ | 91.0 | -49 | -10.9 | -25.5 | -20 | 5.09 |
| Rosmarinic acid | PA | 359.1 $[M-H]^-$ | 160.6 | -49 | -10.9 | -16.5 | -11 | 7.48 |
| Caffeic acid phenethyl ester | PA | 283.09 $[M-H]^-$ | 134.8 | -49 | -10.9 | -29 | -29 | 8.35 |
| 2,3,4-Trihydroxybenzoic acid | PA | 169.0 $[M-H]^-$ | 150.8 | -49 | -10.9 | -16.5 | -16 | 1.15 |
| 2,4-Dihydroxybenzoic Acid | PA | 153.0 $[M-H]^-$ | 109.1 | -49 | -10.9 | -17 | -8 | 2.66 |
| 3-Hydroxybenzoic acid | PA | 137.0 $[M-H]^-$ | 92.8 | -49 | -10.9 | -17 | -27 | 2.74 |
| Coniferyl Alcohol | PA | 179.1 $[M-H]^-$ | 145.6 | -49 | -10.9 | -18 | -5 | 4.5 |
| m-Coumaric acid | PA | 163.04 $[M-H]^-$ | 118.8 | -49 | -10.9 | -18 | -19 | 5.38 |
| 2-Acetylresorcinol | PA | 151.04 $[M-H]^-$ | 134.6 | -49 | -10.9 | -22 | -7 | 5.96 |
| 3,4,5-Trimethoxycinnamic acid | PA | 237.1 $[M-H]^-$ | 103.2 | -49 | -10.9 | -19 | -27 | 7.61 |
| Kaempferol | F | 285.0 $[M-H]^-$ | 197.8 | -56 | -8.5 | -36 | -9 | 8.09 |
| Phloretin | F | 273.1 $[M-H]^-$ | 166.9 | -56 | -8.5 | -24 | -15 | 8 |
| 7,8-Dihydroxyflavone | F | 317.0 $[M-H]^-$ | 178.8 | -56 | -8.5 | -28 | -19 | 7.48 |
| Myricetin | F | 389.1 $[M-H]^-$ | 148.8 | -56 | -8.5 | -38 | -31 | 8.5 |
| Polydatin | F | 301.0 $[M-H]^-$ | 150.3 | -56 | -8.5 | -24 | -5 | 7.88 |
| Quercetin | F | 285.08 $[M-H]^-$ | 184.5 | -56 | -8.5 | -40 | -13 | 8.09 |

| | | | | | | | | |
|---|---|---|---|---|---|---|---|---|
| Acacetin | F | 269.0 [M-H]$^-$ | 194.6 | -56 | -8.5 | -35 | -34 | 8.21 |
| Baicalein | F | 223.1 [M-H]$^-$ | 91.6 | -56 | -8.5 | -38 | -8 | 8.33 |
| 4'-Hydroxychalcone | F | 289.1 [M-H]$^-$ | 252.9 | -56 | -8.5 | -19 | -17 | 7.9 |
| (+)-Catechin (Hydrate) | F | 301.0 [M-H]$^-$ | 150.3 | -56 | -8.5 | -24 | -5 | 7.88 |
| Mangiferin | F | 421.1 [M-H]$^-$ | 300.9 | -56 | -8.5 | -27 | -14 | 5.21 |
| (+)-Taxifolin | F | 303.0 [M-H]$^-$ | 285.0 | -56 | -8.5 | -17 | -20 | 5.97 |
| Diosmetin | F | 299.0 [M-H]$^-$ | 284.0 | -56 | -8.5 | -28 | -21 | 8.16 |
| Morin | F | 301.0 [M-H]$^-$ | 150.5 | -56 | -8.5 | -31 | -32 | 7.85 |
| (-)-Epigallocatechin gallate hydrate | F | 457.1 [M-H]$^-$ | 168.9 | -56 | -8.5 | -16 | -8 | 4.96 |
| Chrysin | F | 253.1 [M-H]$^-$ | 142.8 | -56 | -8.5 | -41 | -41 | 8.42 |
| (+/-)-Naringenin | F | 271.1 [M-H]$^-$ | 118.8 | -56 | -8.5 | -28 | -14 | 7.98 |
| Baicalin | F | 445.1 [M-H]$^-$ | 268.5 | -56 | -8.5 | -27 | -18 | 8.05 |
| Resveratrol | F | 227.0 [M-H]$^-$ | 184.4 | -56 | -8.5 | -22 | -9 | 7.34 |
| Luteolin | F | 285.0 [M-H]$^-$ | 133.0 | -56 | -8.5 | -41 | -37 | 7.3 |
| Hesperidin | F | 609.2 [M-H]$^-$ | 300.7 | -56 | -8.5 | -31 | -31 | 7.63 |
| Fisetin | F | 285.0 [M-H]$^-$ | 134.5 | -56 | -8.5 | -27 | -14 | 7.55 |
| (-)-Epicatechin | F | 289.1 [M-H]$^-$ | 252.8 | -56 | -8.5 | -22 | -22 | 7.75 |
| Oxyresveratrol | F | 243.0 [M-H]$^-$ | 200.3 | -56 | -8.5 | -25 | -6 | 6.64 |
| Apigenin | F | 269.0 [M-H]$^-$ | 116.4 | -56 | -8.5 | -47 | -7 | 8.1 |
| *trans*-Pterostilbene | F | 255.1 [M-H]$^-$ | 239.5 | -56 | -8.5 | -27 | -6 | 8.46 |
| Rutin Hydrate | F | 609.1 [M-H]$^-$ | 299.7 | -56 | -8.5 | -47 | -9 | 7.27 |
| Phloridzin | F | 435.1 [M-H]$^-$ | 272.5 | -56 | -8.5 | -20 | -37 | 7.51 |
| Daidzein | F | 253.1 [M-H]$^-$ | 207.5 | -56 | -8.5 | -35 | -28 | 7.74 |
| Hesperetin | F | 301.1 [M-H]$^-$ | 163.4 | -56 | -8.5 | -39 | -11 | 8.05 |
| Puerarin | F | 415.1 [M-H]$^-$ | 294.9 | -56 | -8.5 | -30 | -12 | 5.09 |
| Isoliquiritigenin | F | 255.1 [M-H]$^-$ | 119.0 | -56 | -8.5 | -27 | -33 | 8.13 |
| Piceatannol | F | 243.06 [M-H]$^-$ | 158.3 | -56 | -8.5 | -26 | -13 | 6.35 |
| Biochanin A | F | 283.1 [M-H]$^-$ | 267.7 | -56 | -8.5 | -26 | -10 | 8.42 |
| Formononetin | F | 267.1 [M-H]$^-$ | 251.5 | -56 | -8.5 | -28 | -4 | 8.21 |
| Diosmin | F | 607.2 [M-H]$^-$ | 298.4 | -56 | -8.5 | -22 | -21 | 7.67 |
| Naringin dihydrochalcone | F | 581.18 [M-H]$^-$ | 273.0 | -56 | -8.5 | -38 | -10 | 7.75 |
| Equol | F | 243.0 [M-H]$^-$ | 200.8 | -56 | -8.5 | -27 | -19 | 6.33 |
| Genistein | F | 269.0 [M-H]$^-$ | 132.7 | -56 | -8.5 | -34 | -16 | 7.98 |

[a]IS, internal standard; PA, Phenolic acid; F, Flavonoid. [b]DP, declustering potential. [c]EP, entrance potential. [d]CE, collision energy. [e]CXP, Collision cell exit potential. [f]RT, retention time.

**Supplementary Table S2**. Linearity and sensitivity data for phenolic acids and flavonoids.

| Metabolite | Metabolite Class[a] | Linearity range (µg/mL) | $r^2$ | LOD[b] | LOQ[c] | Curve equation |
|---|---|---|---|---|---|---|
| o-Anisic Acid | PA | 0.03–0.63 | 0.998 | 0.01 | 0.03 | y = 3E+06x + 44109 |
| 2,3-Dihydroxybenzoic acid | PA | 0.03–5.00 | 0.997 | 0.006 | 0.03 | y = 2E+06x + 189509 |
| Chlorogenic acid | PA | 0.03–5.00 | 0.997 | 0.002 | 0.005 | y = 1E+06x + 87762 |
| Syringic acid | PA | 0.63–10.0 | 0.998 | 0.31 | 0.63 | y = 17964x - 1183.6 |
| p-Coumaric acid | PA | 0.03–2.50 | 0.998 | 0.005 | 0.02 | y = 2E+06x + 122038 |
| m-Hydrocoumaric acid | PA | 0.31–10.0 | 0.999 | 0.03 | 0.31 | y = 943964x + 150482 |
| Ferulic acid | PA | 0.03–10.0 | 0.999 | 0.002 | 0.03 | y = 559017x + 57790 |
| Sinapic acid | PA | 0.03–10.0 | 0.999 | 0.002 | 0.005 | y = 222015x + 3754.9 |
| Aspirin | PA | 0.03–10.0 | 0.998 | 0.002 | 0.007 | y = 405391x - 67347 |
| *trans*-Cinnamic acid | PA | 2.50–10.0 | 0.995 | 1.50 | 2.50 | y = 36651x - 21342 |
| 4-Hydroxybenzoic acid | PA | 0.03–1.25 | 0.998 | 0.006 | 0.03 | y = 2E+06x + 26546 |
| 2,6-Dihydroxybenzoic acid | PA | 0.003–2.50 | 0.997 | 0.001 | 0.003 | y = 4E+06x + 164093 |
| Dihydrocaffeic acid | PA | 0.03–5.00 | 0.999 | 0.002 | 0.007 | y = 538677x + 27979 |
| Caffeic acid | PA | 0.31–2.50 | 0.997 | 0.03 | 0.12 | y = 1E+06x + 153979 |
| Phloretic acid | PA | Not quantifiable in the selected linearity range | | | | |
| Hydroferulic acid | PA | 0.03–5.00 | 0.999 | 0.001 | 0.003 | y = 270948x + 1777.7 |
| Ellagic acid dihydrate | PA | 0.03–1.25 | 0.997 | 0.007 | 0.03 | y = 99817x + 3393 |
| 5-Methoxysalicylic acid | PA | 0.03–5.00 | 0.999 | 0.001 | 0.03 | y = 3E+06x + 126683 |
| Catechol | PA | 0.03–2.50 | 0.996 | 0.001 | 0.004 | y = 1E+06x + 53882 |
| Gentisic acid | PA | 0.003–1.25 | 0.999 | 0.001 | 0.003 | y = 1E+06x + 6905.2 |
| 4-Acetocatechol | PA | 0.03–2.50 | 0.996 | 0.007 | 0.03 | y = 284650x + 23702 |
| 4-Methylcatechol | PA | 0.31–10.0 | 0.999 | 0.03 | 0.14 | y = 7528x + 825.75 |
| 2,6-Dimethoxybenzoic acid | PA | 1.25 – 10.0 | 0.996 | 0.63 | 1.25 | y = 49710x + 38212 |
| Acetylphloroglucinol | PA | 0.03–2.50 | 0.994 | 0.004 | 0.03 | y = 2E+06x + 100214 |
| Salicylic acid | PA | 0.03–10.0 | 0.999 | 0.002 | 0.03 | y = 3E+06x + 402246 |
| trans-2-Hydroxycinnamic acid | PA | 0.03–5.00 | 0.998 | 0.003 | 0.03 | y = 2E+06x + 193871 |
| Caffeic acid dimethyl ether | PA | 0.31–10.0 | 0.998 | 0.03 | 0.08 | y = 42692x - 4380 |
| 3-Methoxyhydrocinnamic acid | PA | 0.31–10.0 | 0.997 | 0.03 | 0.11 | y = 16741x - 2051.4 |
| Gallic acid | PA | 0.03–2.50 | 0.999 | 0.005 | 0.03 | y = 1E+06x + 37728 |
| 3,5-Dihydroxybenzoic acid | PA | 0.31–5.00 | 0.998 | 0.03 | 0.31 | y = 695154x + 108182 |
| Vanillic acid | PA | 0.31–10.0 | 0.999 | 0.03 | 0.01 | y = 25878x - 747.51 |
| Nordihydroguaiaretic Acid | PA | 0.03–2.50 | 0.999 | 0.001 | 0.002 | y = 2E+06x + 21094 |
| Terephthalic acid | PA | 0.03–2.50 | 0.999 | 0.01 | 0.03 | y = 1E+06x + 35990 |
| 4-Acetylresorcinol | PA | 0.03–2.50 | 0.999 | 0.001 | 0.003 | y = 906372x + 17474 |
| Rosmarinic acid | PA | 0.03–5.00 | 0.999 | 0.0005 | 0.001 | y = 1E+06x - 3641.7 |
| Caffeic acid phenethyl ester | PA | 0.03–2.50 | 0.997 | 0.001 | 0.03 | y = 5E+06x + 215577 |
| 2,3,4-Trihydroxybenzoic acid | PA | 0.03–0.60 | 0.999 | 0.001 | 0.003 | y = 874629x + 5590.4 |
| 2,4-Dihydroxybenzoic Acid | PA | 0.03–5.00 | 0.998 | 0.01 | 0.03 | y = 928863x + 55270 |
| 3-Hydroxybenzoic acid | PA | 0.31–5.00 | 0.998 | 0.03 | 0.31 | y = 640365x + 43840 |
| Coniferyl Alcohol | PA | Not quantifiable in the selected linearity range | | | | |
| m-Coumaric acid | PA | 0.03–2.50 | 0.999 | 0.001 | 0.03 | y = 2E+06x + 80336 |
| 2-Acetylresorcinol | PA | 0.03–1.25 | 0.996 | 0.002 | 0.005 | y = 3E+06x + 99822 |
| 3,4,5-Trimethoxycinnamic acid | PA | 0.03–10.0 | 0.998 | 0.0003 | 0.03 | y = 105739x + 13535 |
| Kaempferol | F | 0.31–2.50 | 0.996 | 0.03 | 0.1 | y = 31889x + 313.64 |
| Phloretin | F | 0.03–2.50 | 0.998 | 0.0004 | 0.001 | y = 6E+06x + 181590 |
| Myricetin | F | 0.31–5.00 | 0.995 | 0.05 | 0.15 | y = 669096x - 127523 |
| Polydatin | F | Not quantifiable in the selected linearity range | | | | |
| Quercetin | F | 0.03–2.50 | 0.995 | 0.003 | 0.03 | y = 232809x + 10141 |
| Acacetin | F | 0.03–2.50 | 0.996 | 0.004 | 0.03 | y = 141397x + 7498.9 |
| Baicalein | F | 0.03–5.00 | 0.998 | 0.004 | 0.03 | y = 244740x - 4360.9 |
| 4'-Hydroxychalcone | F | 0.03–2.50 | 0.995 | 0.002 | 0.03 | y = 687940x + 15160 |

| | | | | | | |
|---|---|---|---|---|---|---|
| (+)-Catechin (Hydrate) | F | 0.31–1.25 | 0.997 | 0.03 | 0.06 | y = 98018x + 18549 |
| Mangiferin | F | 0.03–5.00 | 0.999 | 0.001 | 0.003 | y = 1E+06x + 84377 |
| (+)-Taxifolin | F | 0.03–5.00 | 0.999 | 0.002 | 0.005 | y = 2E+06x + 50504 |
| Diosmetin | F | 0.03–1.25 | 0.997 | 0.002 | 0.005 | y = 6E+06x + 112067 |
| Morin | F | 0.03–5.00 | 0.997 | 0.002 | 0.005 | y = 717491x + 68919 |
| (-)-Epigallocatechin gallate hydrate | F | 0.03–10.0 | 0.998 | 0.008 | 0.03 | y = 566243x - 71657 |
| Chrysin | F | 0.03–5.00 | 0.996 | 0.0006 | 0.03 | y = 598355x + 56081 |
| (+/-)-Naringenin | F | 0.03–5.00 | 0.996 | 0.0005 | 0.001 | y = 2E+06x - 15413 |
| Baicalin | F | 0.03–2.50 | 0.997 | 0.001 | 0.003 | y = 1E+06x - 10656 |
| Resveratrol | F | 0.03–10.0 | 0.998 | 0.009 | 0.03 | y = 46605x + 1953.5 |
| Luteolin | F | Not quantifiable in the selected linearity range | | | | |
| Hesperidin | F | 0.03–10.0 | 0.996 | 0.002 | 0.006 | y = 433221x + 1285.5 |
| Fisetin | F | 0.03–5.00 | 0.996 | 0.002 | 0.03 | y = 976215x + 46400 |
| (-)-Epicatechin | F | 0.03–1.25 | 0.996 | 0.01 | 0.03 | y = 141591x + 674.48 |
| Oxyresveratrol | F | 0.03–10.0 | 0.998 | 0.008 | 0.03 | y = 30531x + 1319 |
| Apigenin | F | 0.03–1.25 | 0.994 | 0.001 | 0.005 | y = 763274x - 26083 |
| *trans*-Pterostilbene | F | 0.31–10.0 | 0.996 | 0.007 | 0.03 | y = 52998x - 1462.5 |
| Rutin Hydrate | F | 0.03–5.00 | 0.995 | 0.002 | 0.005 | y = 1E+06x + 15351 |
| Phloridzin | F | 0.03–5.00 | 0.997 | 0.0005 | 0.002 | y = 437141x - 8944 |
| Daidzein | F | 0.03–2.50 | 0.995 | 0.005 | 0.03 | y = 247718x + 22978 |
| Hesperetin | F | 0.03–5.00 | 0.996 | 0.0005 | 0.001 | y = 619584x - 14267 |
| Puerarin | F | 0.03–5.00 | 0.998 | 0.0005 | 0.002 | y = 2E+06x - 1073 |
| Isoliquiritigenin | F | Not quantifiable in the selected linearity range | | | | |
| Piceatannol | F | 0.03–10.0 | 0.997 | 0.009 | 0.03 | y = 105690x + 17925 |
| Biochanin A | F | 0.03–1.25 | 0.993 | 0.0004 | 0.001 | y = 1E+07x + 287361 |
| Formononetin | F | 0.03–0.63 | 0.999 | 0.00001 | 0.003 | y = 2E+07x + 33838 |
| Diosmin | F | 0.03–5.00 | 0.997 | 0.002 | 0.008 | y = 221995x - 1498.1 |
| Naringin dihydrochalcone | F | 0.03–10.0 | 0.999 | 0.004 | 0.03 | y = 418240x + 42251 |
| Equol | F | 0.03–2.50 | 0.997 | 0.002 | 0.005 | y = 727655x + 19681 |
| Genistein | F | 0.03–5.00 | 0.994 | 0.01 | 0.03 | y = 486314x - 46341 |

[a]PA, Phenolic acid; F, Flavonoid. [b]LOD, Limit of detection. [c]LOQ, Limit of quantification.
